# Beyond native sequence recovery: Improved modeling of the sequence-energy landscape of protein structures

**DOI:** 10.64898/2026.01.14.699067

**Authors:** Foster Birnbaum, Amy E. Keating

**Affiliations:** Department of Biology, Massachusetts Institute of Technology, Cambridge, Massachusetts 02139, USA; Computational and Systems Biology, Massachusetts Institute of Technology, Cambridge, Massachusetts 02139, USA; Department of Biological Engineering, Massachusetts Institute of Technology, Cambridge, Massachusetts 02139, USA; Koch Institute for Integrative Cancer Research, Massachusetts Institute of Technology, Cambridge, Massachusetts 02139, USA

## Abstract

Computational protein design using machine learning models has advanced rapidly since the introduction of AlphaFold2. There is now a suite of tools that enable in silico design of proteins with desired structures and properties. Most design workflows require fitting a designed backbone with a sequence that stabilizes it, and many machine learning sequence design models have been proposed. These models are trained to recover the native sequence paired with a known structure, a task known as native sequence recovery (NSR). Here, we demonstrate the limitations of optimizing a sequence design model only for NSR. We show that NSR is often misaligned with more important metrics of model performance: the compatibility of the generated sequence with the desired fold and the ability of the model to predict the energetic effects of mutations. We introduce PottsMPNN, which is trained to generate a Potts energy function consisting of single-residue and residue-pair terms from a protein backbone, and we demonstrate that learning a Potts model reduces NSR but improves sequence generation and energy prediction. To further show that NSR is not the optimal metric, we trained PottsMPNN with noised backbone structures and multiple sequence alignments. In tests on held-out data, NSR decreased, but the quality of the designed sequences and energy predictions improved. By demonstrating the limitations of optimizing for NSR and the effectiveness of alternative strategies for avoiding over-optimizing for NSR, our work provides a new direction for the sequence design field.

## Introduction

Computational protein design holds enormous potential to address outstanding needs in many areas, including human health and sustainability (1). Designing a protein involves selecting a sequence of amino acids that can accomplish a specified function. A common approach to this is to generate a protein backbone structure with desired properties (e.g., shape complementarity to a desired binding target) and then design a sequence that is consistent with that backbone (1). Deep learning methods have rapidly advanced computational protein design and modeling capabilities and can be used for backbone and sequence design (2). Broadly, such techniques involve optimizing a mathematical model defined by a specific functional form and a set of parameters to accomplish a task. The parameters are determined through a training process that involves learning from examples. In the field of protein modeling and design, AlphaFold (versions 1 through 3) is a deep learning method that can predict a protein structure from a protein sequence (3–5), and RFdiffusion is a deep learning method that can generate novel, realistic protein structure backbones (6). Both models were trained using large numbers of experimentally determined or predicted protein structures.

Several deep learning methods have been trained to generate a sequence *s* of amino acids when given a protein backbone *f* : i.e., they have been trained to learn the probability distribution *P_θ_*(*s* | *f*), where *θ* represents the parameters of the model (7–14). Of these models, ProteinMPNN is the most widely used for protein design (9). ProteinMPNN was built on foundational work that established that using a graph neural network (GNN), with residues as graph nodes and model parameters *θ* describing the properties of nodes and relationships between them, facilitates learning sequence-structure relationships (7). In this framework, for a protein of length *L*, *P_θ_*(*s* | *f*) is represented as *L* single-site probability distributions over the 20 standard amino acids that are extracted from the node embeddings in the graph.

Li et al. developed a model, COORDinator, that is based on an alternative framework where *P_θ_*(*s* | *f*) is represented as a Potts model—–a function that decomposes the sequence-energy landscape into a sum of self-energies and pair-energies (8). Potts models and other pairwise interaction analyses, such as double-mutant cycles, have been used to model protein energies for decades because protein folding and stability emerge from networks of cooperative pairwise interactions, such as hydrogen bonds, hydrophobic packing, and salt bridges (15–19). Li et al. and several other groups have showed that learning a pairwise-decomposable energy function can outperform learning single-site probability distributions on some tasks (8, 20–22).

The success of sequence design models can be quantified as the likelihood of the generated sequence folding into the desired structure: i.e., if *s_gen_* := max*_s_ P_θ_*(*s* | *f*) for desired structure *f*, the model is successful if *P* (*f* | *s_gen_*) is high. Tools like Rosetta, which uses a physics- and statistics-based energy function to assess sequence-structure compatibility (23), and AlphaFold are frequently used to assess the quality of the sequence designs by approximating *P* (*f* | *s_gen_*). Many design pipelines generate large numbers of sequences with ProteinMPNN, score them with AlphaFold or Rosetta, and retain only the high-scoring designs (24–27). Such pipelines have generated sequences that adopt desired structures with high experimental success rates.

Because sequence design models learn the probability distribution of residues given a protein backbone, they can also be used to predict the effect of mutations (8, 28–30). This task relies on the learned probability distribution *P_θ_*(*s* | *f*) matching the real sequence-energy landscape for a given structure: i.e., if a model evaluates one sequence to be more likely than another, that should reflect greater protein stability. Given a structure *f* and two sequences *s_wt_* and *s_mut_*, a sequence design model can predict the energy difference as ΔΔ*G_pred_*(*s_wt_, s_mut_*) = log *P_θ_*(*s_mut_* | *f*) − log *P_θ_*(*s_wt_* | *f*), which can be compared to an experimentally observed ΔΔ*G_expt_*(*s_wt_, s_mut_*). Because sequence design models are trained to predict the probability of a sequence given a structure, they should be evaluated using physical properties (e.g., folding and binding energies) that are closely tied to structural features as opposed to molecular or cellular functions (e.g., effects on protein expression or cell growth) that are indirect readouts of such thermodynamic features (31).

The two use cases described above provide two optimization objectives for sequence design models: (1) sequence-structure self-consistency, quantified by *P* (*f* | *s_gen_*), which assesses the compatibility of the designed sequence with the desired structure, and (2) mutation-effect prediction, quantified by the correlation *Pearson*(ΔΔ*G_expt_,* ΔΔ*G_pred_*) between experimentally observed and predicted mutation energies. Unfortunately, there are impediments to training on either objective. The sequence-structure self-consistency scores are either non-differentiable with respect to sequence (Rosetta) or slow to compute and difficult to optimize in sequence space (AlphaFold). And although energy prediction is an effective objective on which to fine-tune the weights of a previously trained model (29, 30), there is insufficient experimental energy data to train a generalizable model from scratch.

Because of these limitations, sequence design models are trained on the task of native sequence recovery (NSR). Given a pairing of backbone structure, *f_nat_*, and native sequence, *s_nat_*, from the Protein Data Bank (PDB) (32) or AlphaFoldDB (28, 33, 34), the task is to maximize *P_θ_*(*s_nat_* | *f_nat_*) so that *s_gen_* ≈ *s_nat_*. Assuming that *P* (*f_nat_* | *s_nat_*) is high and that *P* (*f_nat_* | *s_gen_*) approaches *P* (*f_nat_* | *s_nat_*) as *s_gen_* approaches *s_nat_*, training on NSR optimizes for *P* (*f* | *s_gen_*). Due to the pairing of *f_nat_* and *s_nat_* and the relationship between sequence similarity and structural similarity for native proteins (35), both assumptions are reasonable. Also, training on NSR is empirically effective. The reference energies for amino acids in the Rosetta energy function were set by optimizing for NSR (23, 36), and Rosetta has been successfully used to design many proteins (37). ProteinMPNN was trained on NSR and can design sequences that fold to intended structures and can be fine-tuned to predict energies with state-of-the-art performance (29, 30).

However, there are several reasons why optimizing for continual improvement in NSR may fail to produce corresponding improvements in sequence-structure self-consistency and energy prediction. A fundamental challenge is that the relationship between protein sequence and structure is many-to-many, not one-to-one. For any given fold *f_nat_*, *s_nat_* is not the only sequence that will adopt that fold; many proteins that share only 40% sequence identity adopt highly similar structures (38, 39). Furthermore, the native sequence is usually not the sequence that maximizes the stability of a structure. Deep mutational scanning experiments show that although most mutations in native proteins are energetically unfavorable, many positions have at least one favorable substitution relative to wildtype (40–42). Furthermore, a protein sequence *s_nat_* does not adopt a single structure: a particular concern is that the structures of proteins are over-specified by the atomic coordinates deposited in the PDB, which capture a single conformation that lies in a local optimum of the force field used for structure refinement. Optimizing for similarity to *s_nat_* ignores sequence-structure degeneracy and the fact that evolution may select for sequences that are good enough to function and not necessarily optimal in any way. Continuously maximizing *P_θ_*(*s_nat_* | *f_nat_*) may therefore result in an unrealistic model of the sequence-energy landscape.

We benchmarked state-of-the-art sequence design models on sequence-structure self-consistency and energy prediction. We found that Frame2Seq and ProteinMPNN achieved the best performance, even though they generate sequences with lower NSR compared to other models. We then tested several ways of altering the training task to better reflect principles underlying protein sequence-structure relationships. First, we developed PottsMPNN, a sequence design model that is trained to learn a Potts model and, simultaneously, single-site amino-acid probability distributions. PottsMPNN achieved lower NSR but outperformed all models on sequence-structure self-consistency and energy prediction. Second, consistent with an observation by Dauparas et al., we found that training with coordinate noise improves sequence-structure self-consistency (9). We demonstrate that introducing noise during training also improves energy prediction and that training with noise is beneficial because it prevents overfitting to NSR during training. Third, and most importantly for reducing reliance on NSR, we trained to recover information from multiple sequence alignments (MSAs) to directly provide the model with examples of related sequences that adopt a shared fold. Doing so improved both sequence-structure self-consistency and energy prediction. Our results demonstrate that NSR is not the right objective for state-of-the-art sequence design models and that optimizing for more biologically appropriate objectives increases model performance.

## Results

### PottsMPNN architecture

PottsMPNN is a GNN that learns to encode a protein backbone as a graph, extract a Potts model from the edge embeddings in the graph, and autoregressively decode the node embeddings to predict single-site probability distributions (Fig. 1, see Methods for details). The autoregressive decoder is convenient for sequence generation, and the Potts model is used to score sequence variants, such as those resulting from point mutations. PottsMPNN integrates two models: COORDinator (8) and ProteinMPNN (9). PottsMPNN uses the same featurization and GNN encoder as ProteinMPNN. After the encoder, a single neural network layer applies a linear function to reshape the edge embeddings into the form of a Potts model. Critically, the Potts model is supervised using the negative log composite pseudo-likelihood loss developed for COORDinator, which rewards the model for assigning a high probability to the native pair of residues at a given pair of positions (graph edge loss L*_E_*). As observed by Li et al., the Potts model learned using this loss function can predict energies effectively, but the sequences derived from using Markov chain Monte Carlo (MCMC) to sample from the Potts model are often low complexity, and running MCMC sampling takes a long time (8). Accordingly, PottsMPNN incorporates the autoregressive GNN decoder from ProteinMPNN to generate single-site probability distributions at each graph node, which are supervised to reward the model for assigning a high probability to the native residues at each site (graph node loss L*_V_*). The model is trained using a composite loss that is the sum of L*_E_* and L*_V_*, equally weighted.

**Fig. 1.**
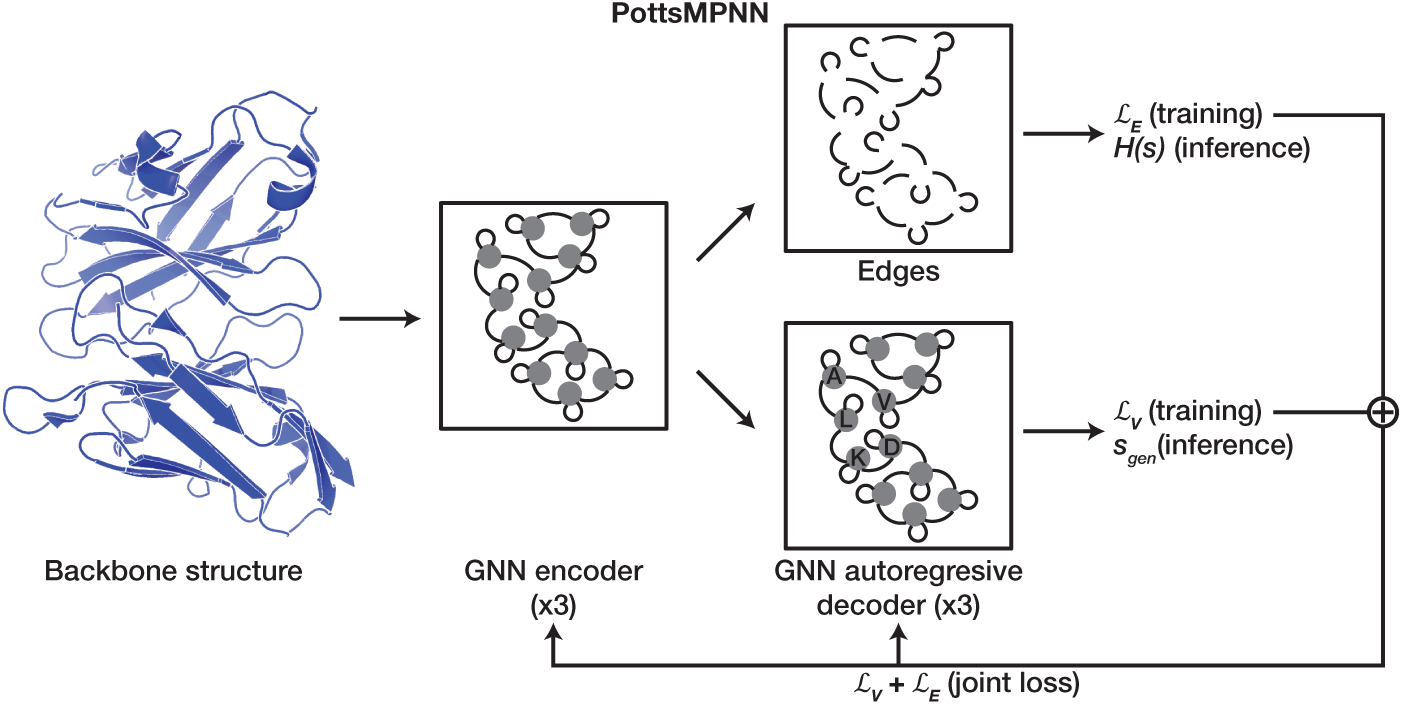
Overview of the PottsMPNN architecture. The model input is a protein structure backbone that is used to define a *k*-NN graph. The nodes and edges of the graph are encoded using a message-passing neural network. The edges, including self-edges, are supervised to learn single and pairwise residue interaction energies in the form of a Potts model (*H*(*s*)), which is used to compute structure energies. The Potts model is supervised using a negative log composite pseudo-likelihood loss (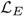) that maximizes the probabilities of native-residue pairs. The nodes are decoded autoregressively to generate single-site amino-acid probabilities and are supervised by a negative log-likelihood loss (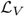) that maximizes native sequence recovery; during inference, the nodes are used to generate a sequence (*s_gen_*). See Methods for details.

### Model benchmarking

We compared PottsMPNN with six other sequence design models: ProteinMPNN (9), MapDiff (14), Frame2Seq (10), UniIF (13), KW-Design (12), and PiFold (11). We tested ProteinMPNN because it provided the model architecture that we used to train PottsMPNN and because it is the most widely used model for protein design. We selected the other five models to include those with among the highest reported NSR values (MapDiff and UniIF), those with intermediate NSR values (KW-Design and PiFold), and those shown to be experimentally successful even at low NSR (Frame2Seq). Because most models were trained on the non-redundant CATH 4.2 dataset (7) that consists only of single-chain proteins, we trained PottsMPNN and retrained ProteinMPNN on that dataset. Because ProteinMPNN was originally trained on the larger PDB-clust dataset (9) that includes multichain proteins, we also trained versions of PottsMPNN on PDB-clust. See Datasets for details on training and testing data. Sequence-structure self-consistency was evaluated on the CATH 4.2 or PDB-clust test sets in two ways. First, we predicted structures of generated sequences with AlphaFold2 and calculated the TM-score (43) between predicted and native structures. (TM-score is a metric of structural similarity that ranges from 0 to 1, with 0 indicating no structural similarity and 1 indicating structural identity.) Second, we calculated length-normalized Rosetta energies for proteins constructed by fitting generated sequences onto the native structures. See Sequence-structure self-consistency for details. Energy prediction was evaluated on three datasets reporting mutational effects on protein stability (see Energy prediction for details) (41, 42, 44). Figure 2 shows the results of benchmarking sequence design models trained without noise using the CATH 4.2 dataset on NSR, sequence-structure self-consistency, and energy prediction. As shown in Fig. 2A, the NSR of ProteinMPNN and PottsMPNN are very low relative to other models: the best model, MapDiff, achieves ~58% NSR, while PottsMPNN achieves only ~44% NSR. However, as shown in Figs. 2B-C, the trend is very different for sequence-structure self-consistency. PottsMPNN achieves the highest sequence-structure self-consistency: PottsMPNN sequences folded by AlphaFold2 have a significantly higher average TM-score to the native structures compared to the sequences generated by all other models (Fig. 2B), and PottsMPNN sequences threaded onto the native structure have significantly better Rosetta scores than the sequences generated by all other models except Frame2Seq (Fig. 2C). The improved sequence-structure self-consistency of PottsMPNN sequences is also apparent when examining the fraction of sequences with TM-scores above various thresholds (SI Appendix, Fig. S1) and the confidence of the predicted structures (SI Appendix, Fig. S2). For energy prediction, PottsMPNN outperforms the other models by a significant margin (Fig. 2D). A case study of 2KXD (45, 46), a protein in the Megascale protein stability test set, shows that at all ranges of ΔΔ*G_expt_* values PottsMPNN predictions are more correlated with experimental values than ProteinMPNN predictions are (SI Appendix, Figs. S3A, S3B). Coloring the structure by per-site Pearson r values shows that PottsMPNN is especially good at predicting the effects of mutations in the structured core of the protein (SI Appendix, Figs. S3C, S3D).

**Fig. 2.**
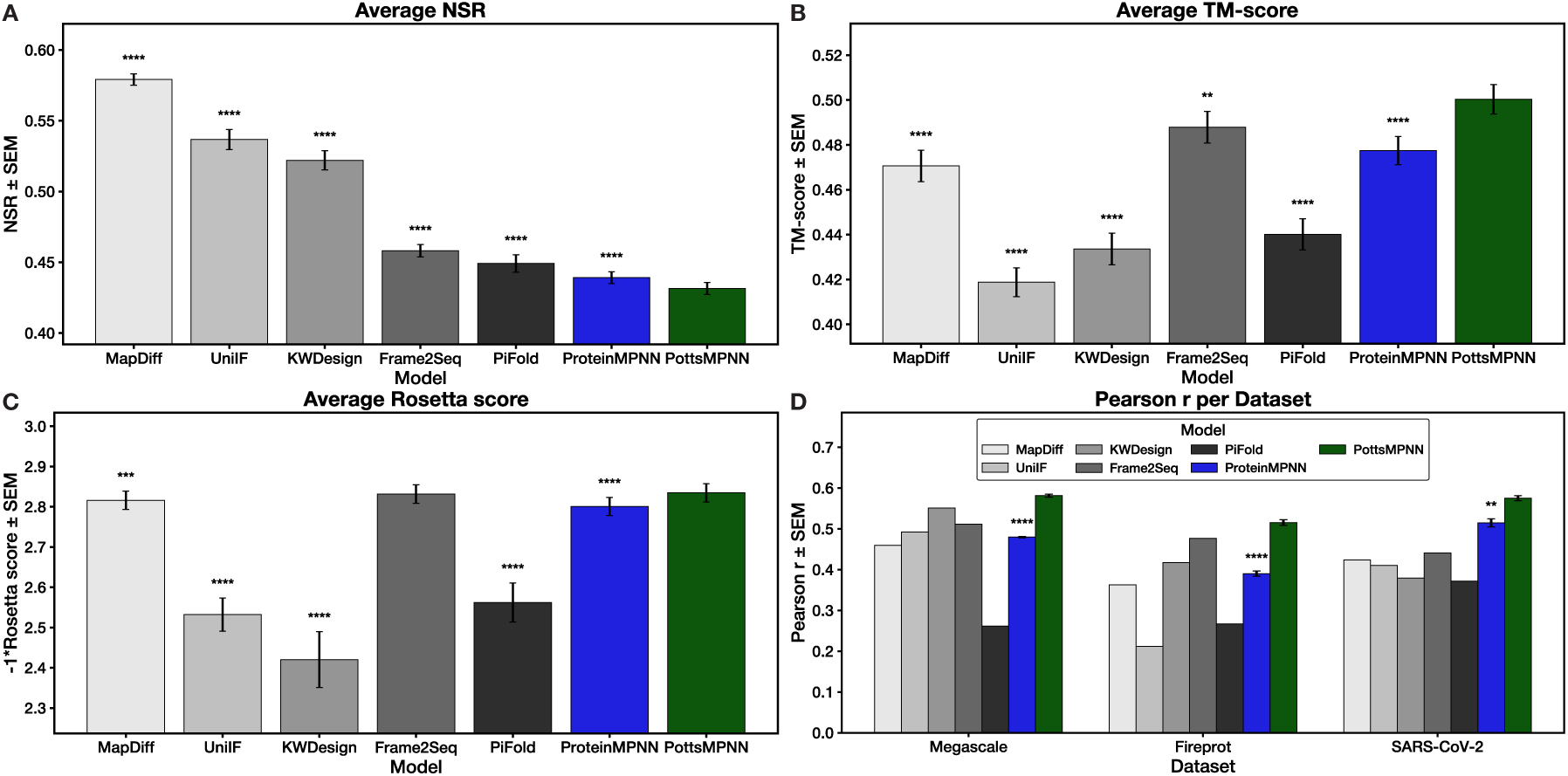
NSR does not correlate with performance on sequence-structure self-consistency or energy prediction for models trained and tested on the CATH 4.2 dataset. (A) NSR results for each model. (B) TM-scores between native structures and AlphaFold2 predicted structures for sequences generated using each model. (C) Rosetta scores (multiplied by −1) after threading generated sequences onto the native backbone and relaxing for each model. (D) Pearson r for using each model to predict the effect of single-site mutations on protein stability for three datasets. For (A) – (C), error bars show SEM over the proteins in the test set after averaging results over three retrained model replicates; for (D), error bars show SEM over three retrained model replicates (where available). Stars indicate statistical significance relative to PottsMPNN, assessed using a two-tailed paired t-test over per-protein values for (A) – (C) and a two-tailed unpaired t-test over average Pearson r values for (D) (* p *<* 0.05, ** p *<* 0.01, *** p *<* 0.001, **** p *<* 0.0001; for (A) – (C), no star indicates non-significance).

### Local sequence optimization using energy predictions

Given that the Potts model performs well when predicting the effects of single-site mutations, we hypothesized that it could improve the sequences generated autoregressively using the single-site probability distributions. To test this, we visited each position in the sequence and used the Potts model to find favorable substitutions (see Sequence generation for details). We tracked how sequence-structure self-consistency changed during the local optimization process, and we found that both TM-scores and Rosetta scores consistently improved, with notable alignment between the change in energies predicted by the Potts model and the change in Rosetta scores (SI Appendix, Fig. S4). Energies predicted using single-site probabilities did not reflect that the sequences improve in quality as the local optimization progresses (SI Appendix, Fig. S4). Iterative optimization to convergence did not result in continued improvement in sequence quality after each site had been visited once (SI Appendix, Fig. S5). Because the local optimization requires scoring every possible mutation at all positions in the sequence, it requires more inference time, but sequence generation remains fast at 0.32 ± 0.16 seconds per sequence (SI Appendix, Fig. S6).

### The importance of the Potts model

We explored why PottsMPNN is superior to ProteinMPNN for energy prediction. PottsMPNN learns a Potts model from the edges in the graph embedding of the protein—by supervising joint probability distributions at pairs of sites—and simultaneously learns single-site probability distributions. In contrast, ProteinMPNN only learns single-site probability distributions. To test if the Potts model learned by PottsMPNN is critical to its performance, we compared the sequence scoring ability of the Potts model from PottsMPNN with the sequence scoring ability of the single-site probability distributions from PottsMPNN. Although these two functions are learned simultaneously during training, subject to a joint loss function, using the single-site probabilities to predict energies resulted in significantly worse performance than using the Potts model; using the single-site probabilities from PottsMPNN performs equivalently to using the single-site probabilities from ProteinMPNN (SI Appendix, Fig. S7A). Consistent with this, when we used the single-site probabilities for local optimization of sequences generated using the autoregressive decoder, we observed significantly lower NSR and lower sequence-structure self-consistency scores than we did when using the Potts model for local optimization (SI Appendix, Figs. S7B-S7D). Indeed, the TM-scores for single-site locally optimized sequences were significantly worse compared with the original sequences (SI Appendix, Fig. S7C). The Potts model energies reflected that the sequences do not improve in quality as the single-site local optimization progresses (SI Appendix, Fig. S8). These tests establish the utility of the Potts function as an output of the model.

To test whether supervising residue pairs is critical to learning an informative Potts model, we trained two versions of PottsMPNN: PottsOnlyMPNN and PottsSingleMPNN (SI Appendix, Fig. S9; see Loss functions and optimization for details). These models are trained only to learn a Potts model from the graph edges, with no single-site probability distribution track. PottsOnlyMPNN is trained with the normal negative log composite pseudo-likelihood loss that supervises residue identities at pairs of sites; it is similar to COORDinator (8). PottsSingleMPNN, in contrast, learns a Potts model but supervises only single-residue identities. Specifically, PottsSingleMPNN is supervised on the single-site negative log likelihood loss when each single-site probability distribution is calculated from the Potts model by summing the self-energies at that site and all pair-energies conditioned on the native residue identities at all other sites. PottsOnlyMPNN performs similarly to PottsMPNN at energy prediction, but PottsSingleMPNN performs significantly worse, supporting the importance of supervising residue pairs during training (SI Appendix, Fig. S7).

We also tested whether certain pairwise interactions in the Potts model are more important than others by summing over different numbers of residue pairs at inference time. Progressively removing residue pairs significantly reduced energy prediction performance (SI Appendix, Fig. S10). Removing interactions between near neighbors had a greater effect than removing interactions between far neighbors for most tests.

### Adding noise during training

We tested the effect of adding increasing amounts of noise to the input structures during training. For these experiments, models were trained on the larger PDB-clust dataset. Following Dauparas et al., noise was added by perturbing the position of each atom by sampling a displacement in each coordinate axis independently from a Gaussian distribution of a set standard deviation, *σ*, which we refer to as the noise level (9). As the noise level added during training increased, models generated sequences with lower NSR but significantly higher sequence-structure self-consistency scores (up to *σ* = 0.3 Å), and they performed significantly better at energy prediction on the Megascale test set (up to *σ* = 0.2 Å) (Fig. 3A-C). To investigate why adding noise during training improves performance, we compared the training and validation loss curves and sequence-structure self-consistency and energy prediction performance at selected epochs of training for two models: one trained without noise (Fig. 3D-F) and one trained with *σ* = 0.2 Å (Fig. 3G-I). The model trained without noise exhibits signs of overfitting to NSR: the generated sequences do not improve according to both sequence-structure self-consistency methods, while the NSR training and validation losses continue to decrease. However, the model trained with noise exhibits the opposite behavior, as sequence-structure self-consistency generally improves as it is trained. Also, the model trained with noise shows greater improvement in energy prediction over the course of training. The train loss for the noise model is higher than the validation loss because noise is only added during training.

**Fig. 3.**
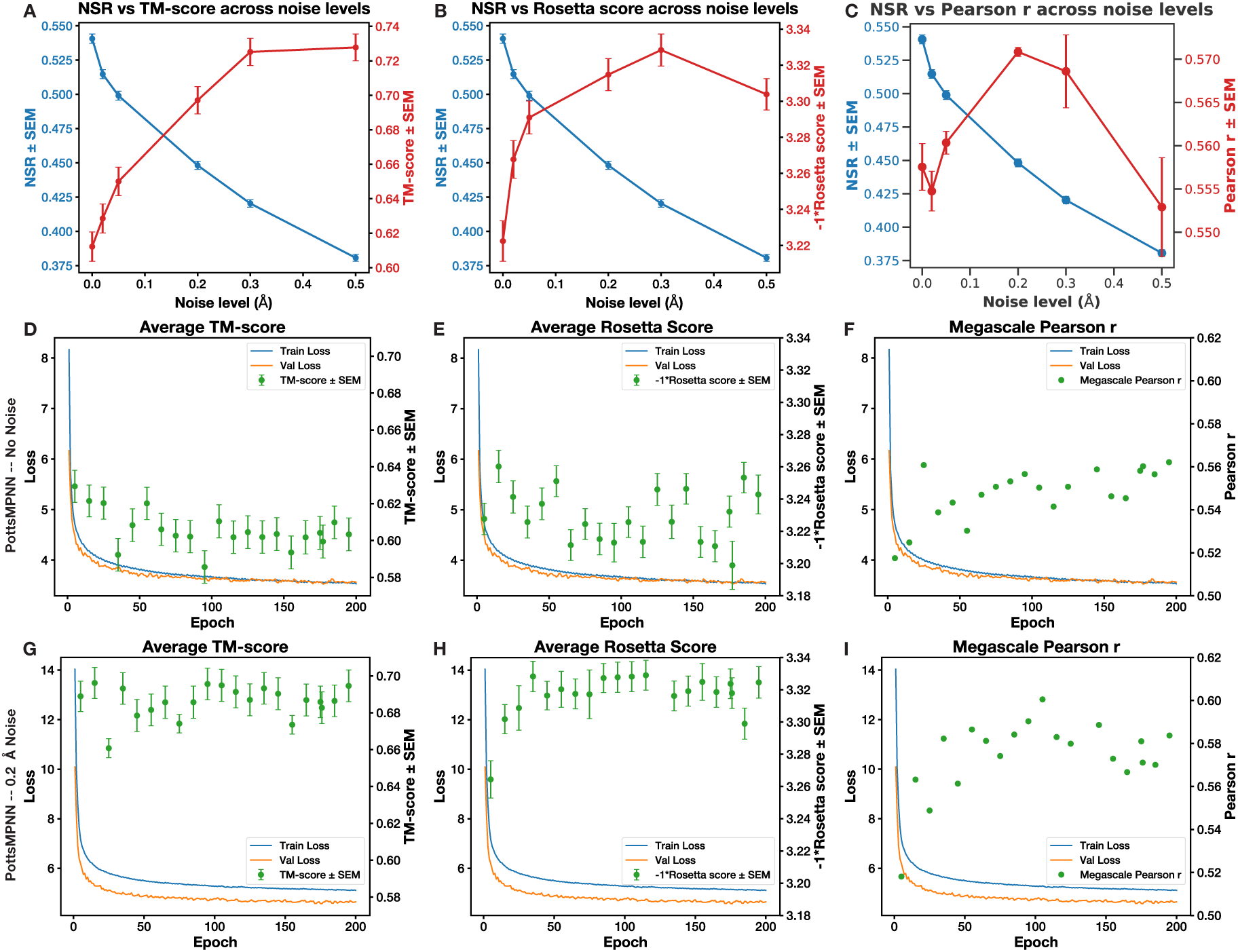
Training PottsMPNN on the PDB-clust dataset with noise improves performance and prevents overfitting to NSR. (A) NSR decreases, whereas the TM-score between the native structure and AlphaFold2 structure predicted from generated sequences improves with increasing noise. (B) Rosetta scores from modeling generated sequences on native backbones improve with increasing noise up to 0.3 Å. (C) Pearson r for predicting the Megascale single-residue mutation energies improves with increasing noise up to 0.2 Å. For (A) – (B), error bars represent SEM over the proteins in the test set after averaging results over at least two retrained model replicates; for (C), error bars show SEM over retrained model replicates. (D) – (F) TM-score (D), Rosetta score (E), and Megascale Pearson r values (F) for various epoch checkpoints when PottsMPNN was trained without noise. (G) – (I) TM-score (G), Rosetta score (H), and Megascale Pearson r values (I) for various epoch checkpoints when PottsMPNN was trained with 0.2 Å noise. For (D), (E), (G), and (H), error bars show SEM over the proteins in the PDB-clust test set.

### Training using MSAs

Finally, we tested the effect of modifying the loss function by averaging the loss over sequences sampled from an MSA (see Loss functions and optimization for details). We filtered each MSA to increase the probability that all sequences in the filtered MSA adopt the same fold while maintaining sequence diversity (see Datasets for details). Our filtered MSAs have a median depth of 147 sequences (interquartile range of 416) for CATH 4.2 and 133 sequences (interquartile range of 502) for PDB-clust (SI Appendix, Fig. S11). We compared models trained on the CATH 4.2 dataset with and without noise and with and without MSAs. Training on MSAs significantly reduced NSR (Fig. 4A) but significantly increased sequence-structure self-consistency (Fig. 4B-C). There is a positive, albeit modest, effect of using MSAs on energy prediction performance (Fig. 4D). The trends in performance are consistent, regardless of the amount of noise, so the best model is trained with noise and MSAs. Using multiple sequences from the MSA during each training iteration is key: when a single random sequence is chosen for each protein for each iteration, performance does not improve (SI Appendix, Fig. S12).

**Fig. 4.**
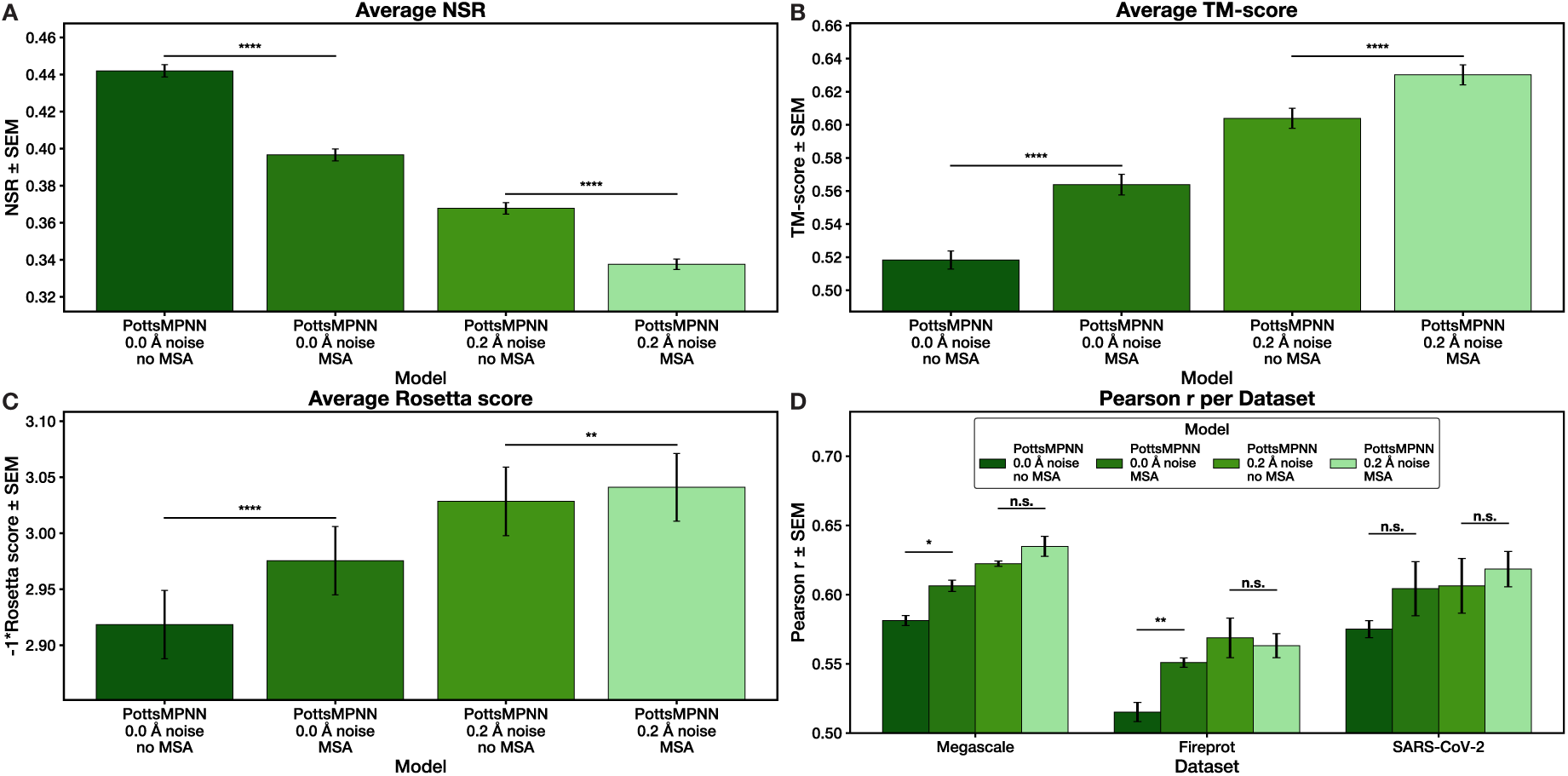
Training on the CATH-4.2 dataset with an MSA loss function improves performance. (A) NSR for models trained with and without noise and with and without an MSA loss function. (B) TM-scores between native structures and AlphaFold2 structures predicted for sequences generated using each model. (C) Rosetta scores after modeling generated sequences on the native backbone. (D) Pearson r for using each model to predict the effect of single-site mutations on protein stability for three datasets. For (A) – (C), error bars show SEM over the proteins in the test set after averaging results over at least two retrained model replicates; for (D), error bars show SEM over at least two retrained model replicates. Stars indicate statistical significance comparing the model trained without MSAs to the model trained with MSAs in each condition, assessed using a two-tailed paired t-test over per-protein values for (A) – (C) and a two-tailed unpaired t-test over average Pearson r values for (D) (* p *<* 0.05, ** p *<* 0.01, *** p *<* 0.001, **** p *<* 0.0001; n.s. indicates non-significance).

We compared PottsMPNN and ProteinMPNN trained on the larger PDB-clust dataset with and without noise and with and without MSAs. Both models improve when training with noise and MSAs, and PottsMPNN performs better in both conditions (SI Appendix, Fig. S13). To investigate the difference between training on the CATH 4.2 and PDB-clust datasets, we compared PottsMPNN models trained on both datasets with and without noise and with and without MSAs. When evaluated on a subset of the PDB-clust test set made non-redundant with the CATH 4.2 train set, the PDB-clust models perform better on sequence-structure self-consistency and energy prediction (SI Appendix, Fig. S14).

Because PottsMPNN relies on a graph representation of the protein backbone with a fixed number of nodes and edges, and because an assumption of the MSA approach is that *P* (*f_nat_* | *s*) is high for all *s* ∈ MSA, sequences in the MSA with gaps or insertions present a difficulty to the model. For a gap, we masked that position and all pairwise interactions that involve that position from affecting the loss calculation. We ignored insertions. We tested various filtering hyperparameters: the minimum sequence identity to the native sequence, the maximum gap percentage, and the maximum insertion percentage. The minimum sequence identity hyperparameter has the largest effect on performance, as models trained with a 70% minimum perform worse across all tasks compared to models trained with a 50% minimum and the same noise level (SI Appendix, Fig. S15). Changing the other hyperparameters did not have large effects, and while no set of hyperparameters resulted in the best performance across all tests, using a 50% sequence identity minimum, a 20% gap maximum, and a 20% insertion maximum resulted in strong performance across all tests (SI Appendix, Fig. S15).

## Discussion

State-of-the-art sequence design models are trained on NSR. By training to maximize *P_θ_*(*s_nat_* | *f_nat_*), they rely on several assumptions, most notably that maximizing the similarity between *s_gen_* and *s_nat_* also maximizes *P* (*f_nat_* | *s_gen_*). Our results show that models trained on NSR–—especially ProteinMPNN and Frame2Seq, which achieve NSR values that are low compared to other models–—can achieve strong performance on sequence-structure self-consistency and decent performance on energy prediction. The high quality of Frame2Seq sequences is consistent with the finding of Akpinaroglu et al. that Frame2Seq can generate sequences with 0% NSR that fold to stable structures with secondary structures consistent with the input backbone structures (10). In general, however, our results demonstrate that optimizing solely for NSR is insufficient to drive further model improvement: compared to ProteinMPNN and Frame2Seq, models like MapDiff and UniIF achieve significantly higher NSR but perform poorly on the sequence-structure self-consistency and energy prediction tests (Fig. 2). We demonstrate the effectiveness of three strategies for improving the training objective, motivated by better aligning the sequence design models with established principles of protein sequence-structure relationships.

First, learning a Potts model *H_θ,E_*(*s* | *f_nat_*) improves model performance (Figs. 2; SI Appendix, Figs. S1, S2, S3). The basis for our work on PottsMPNN is the COORDinator model developed by Li et al., which learns a Potts model and performs well on energy prediction tasks compared to contemporary models (8). We establish that the COORDinator framework can be improved by incorporating an additional loss based on single-site probabilities, L*_V_*, and using single-site probabilities for auto-regressive sequence design, as is done by ProteinMPNN. PottsMPNN, a model trained with a joint objective, generates high-quality sequences as assessed by sequence-structure self-consistency. Sequence quality can be further improved by local optimization using the Potts model energies, indicating that Potts model energies are predictive of sequence quality in the local space around the initial sequence (SI Appendix, Fig. S4). However, continuing the Potts model optimization until convergence did not improve sequence quality, indicating that the Potts model may not be as informative in the global sequence space (SI Appendix, Fig. S5).

Performing local optimization using the Potts model allows PottsMPNN to use the high-quality energetics information in the Potts model without resulting in low complexity sequences, as Li et al. observed when using MCMC to find a global minimum in the Potts model sequence-energy landscape. Another way to avoid low complexity sequences is to apply a complexity penalty term while sampling. Li et al., Ingraham et al., and Shuai et al. demonstrated this approach can work, but we found that even penalizing low-complexity sequences while sampling does not perform as well as and is much slower than autoregressive generation, even when this is followed by local energy optimization (8, 22, 47).

We postulated that the ability of PottsMPNN to optimize sequences and compute energies derives from the explicit residue-pair terms in the Potts model. Learning pairwise interactions allows PottsMPNN to explicitly capture physical constraints (e.g., electrostatic repulsion between proximal residues with the same charge) in a way that a model that only learns single-site probability distributions must learn implicitly through the message passing between sites that occurs in the decoder. This could allow PottsMPNN to capture more complex physical constraints. Although protein energetics involve many-body effects, approximating protein energetics using only self-and pair-energies has proven effective (48, 49).

We performed experiments that withheld all (SI Appendix, Figs. S7, S8) or partial (SI Appendix, Fig. S10) residue-pair information from the model during inference, and we compared models trained with a single-residue loss function to those trained with residue-pair loss functions (SI Appendix, Figs. S7, S9). In all cases, the performance of PottsMPNN degraded when the residue-pair information in the Potts model was limited. The near- and far-neighbor ablation series suggest that model performance starts saturating when each residue has information about its nearest 24 or its furthest 32 of its 48 neighbors, which is consistent with previous observations on the effect of changing the number of nearest neighbors in the *k*-NN graph on the performance of ProteinMPNN (9).

Second, we found that adding noise improves model performance (Fig. 3A-C). This idea was introduced in the context of ProteinMPNN, and the most used version of ProteinMPNN was trained with 0.2 Å of noise. Dauparas et al. suggested that training with noise is beneficial because most training structures were resolved using X-ray crystallography, and structure refinement may encode sequence identity in the backbone coordinates in ways that can be learned but that do not generalize (i.e., in ways that models can memorize) (9). We found evidence consistent with the hypothesis that training with noise provides a regularizing effect that prevents the model from overoptimizing to NSR by measuring trends in sequence-structure self-consistency and energy prediction over the duration of training with and without noise (Fig. 3D-I).

Third, training using an MSA recovery objective instead of an NSR objective improves performance (Figs. 4; SI Appendix, Fig. S13). Because many evolutionarily related proteins adopt closely related structures, we reasoned that including information about homologs would improve the ability of the model to capture meaningful sequence-structure relationships. MSA-based training rewards models that recognize that many sequences can adopt a given structure and prevents over-optimization to a native sequence that may not be optimal in any meaningful sense. We found that averaging the loss over many sequences from the MSA during each iteration is essential: sampling a single sequence from the MSA each iteration does not improve performance (SI Appendix, Fig. S12). We also found that model performance decreases if the MSAs are filtered too stringently so that the sequences in the MSA do not provide diversity (SI Appendix, Fig. S15). We did not explore other ways to improve the use of information in the MSA, such as weighting sequences by their similarity to other sequences in the MSA to increase the effective sequence diversity seen by the model, so we expect that work in this area could result in further performance improvements.

We trained models with and without noise and with and without MSAs on two different datasets, CATH 4.2 and PDB-clust, and observed consistent performance trends. CATH 4.2 is substantially smaller and has a more rigorous, structure-based train-test split, making it ideal for experimenting with different model architectures and optimization objectives. However, models trained on PDB-clust tend to perform better on shared held-out test sets (SI Appendix, Fig. S14). This is likely due to the increased structural diversity in the PDB-clust dataset. Because energy data are not used in training, limiting the possibility of data leakage, we recommend using large structure sets when training models for energy prediction.

Learning residue-pair energies, introducing coordinate noise, and training on homologs of a target sequence do not improve NSR. Nevertheless, these modifications result in a model that is superior at the two tasks that sequence design models are most used for: generating a sequence that folds into a desired structure and predicting the energies of mutations given a structure. Thus, our results demonstrate that NSR should not be the primary metric of success for sequence design models.

There is substantial room for further progress on moving sequence design models away from optimizing for NSR. Several groups have demonstrated that sequence design models can be fine-tuned on energy data, but the generalization of such fine-tuning remains underexplored (29, 30). Also, optimizing only in sequence space leaves the model unaware of potential off-target folding: i.e., the model does not directly learn to minimize *P* (*f_off_* | *s_gen_*), for off-target fold *f_off_*. Pacesa et al. developed BindCraft to address this by iteratively updating *s_gen_* according to a gradient derived from AlphaFold predictions to directly optimize *s_gen_* for *P* (*f_on_* | *s_gen_*) (50). BindCraft generates sequences with high experimental success rates—–after redesigning part of the generated sequence with ProteinMPNN—–indicating that sequence design models that are supervised in structure space instead of sequence space are promising. Accordingly, future work should explore the idea of training or fine-tuning a sequence design model directly on sequence-structure self-consistency or other desired tasks.

## Methods

### Potts model

A Potts model is a function *H*(*s*) that decomposes the sequence-energy landscape for a sequence *s* of length *n* into a sum of self-energies and pair-energies: 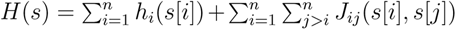, where 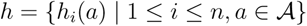 is a lookup table for self-energies, 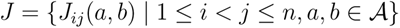 is a lookup table for pair-energies, and 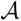 is the set of all 20 amino acids. Several machine learning sequence design models have successfully parameterized a Potts model with learned weights *θ*: they learn *H_θ_*(*s* | *f_nat_*) by maximizing 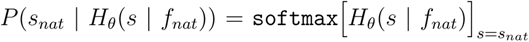 (see Loss function for details) (8, 47).

### Model architecture

We use the same GNN architecture and hyperparameters as ProteinMPNN (9). In brief, the protein backbone is encoded as a *k*-nearest neighbors (*k*-NN) graph, with *k* = 48. Nodes *V* in the graph are residues and are initialized with null vectors. Edges *E* are interactions between residues. For each pair of residues, the interaction representation is initialized using radial basis functions to parameterize the interatomic distances for all 25 pairs of backbone heavy atoms, including virtual C*β* atoms. The node and edge embeddings are then updated using a three-layer message passing neural network (MPNN) encoder. Finally, the node embeddings are updated using a three-layer MPNN autoregressive decoder that selects residues incrementally, conditioning on previously selected residues. The encoder and decoder use a hidden dimensionality of 128. The final edge embeddings (including self-edges) are converted to the Potts model *H*_(_*_θ,E_*_)_(*s* | *f*) using a single linear layer. (Fig. 1)

### Datasets

To benchmark against other models, we used the CATH 4.2 dataset, which consists of 19,700 single-chain structures split 80/10/10 on CATH protein structure classification codes (7, 51). We also trained on the larger set of structures that was used to train ProteinMPNN. This dataset was created by clustering chains from the PDB at 30% sequence identity, creating 25,361 clusters split 90/5/5 such that no chain in the training set is in a complex with chains in the validation or test clusters (9, 32). When using this dataset during training, following Dauparas et al., a new member from each cluster was randomly selected every two epochs (9). We refer to this dataset as the PDB-clust dataset. Because training on CATH 4.2 is much faster, we generally experimented with models trained on that dataset. We note when models are trained on the PDB-clust dataset.

MSAs for 140,000 unique protein chains were downloaded from the OpenFold OpenProteinSet database (52). These MSAs were generated using HHblits (-n3)(53) and JackHMMER (54) and searching against the MGnify (55), BFD (4), UniRef90 (56, 57), and UniClust30 (58) databases. For each complex in the PDB-clust dataset, we create a paired MSA from the OpenFold chain MSAs following the AlphaFold-Multimer species-pairing methodology (59, 60). For each chain in the CATH 4.2 dataset, we generated MSAs using the ColabFold MSA server (61). We filtered sequences in all MSAs using three metrics: a minimum sequence identity to the native sequence, a maximum percentage of insertions, and a maximum percentage of deletions. We experimented with various filters, and empirically the best performance on our tests resulted from using 50% minimum sequence identity, 20% maximum insertions, and 20% maximum deletions.

### Structural noise

To add noise during training, we moved each backbone atom by independently sampling a displacement in all three axes from a Gaussian distribution with mean 0 Å and standard deviation *σ Å*, as done by Dauparas et al. (9). We tested *σ* ∈ [0, 0.02, 0.05, 0.2, 0.3, 0.5].

### Loss functions and optimization

We trained PottsMPNN using an equally weighted composite of two loss functions. First, we supervised the nodes using the negative log likelihood loss used to train ProteinMPNN (9), shown in Equation 1:

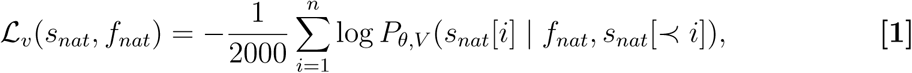

where *n* is the sequence length, *P_θ,V_* (*s*[*i*] | *f_nat_, s_nat_*[≺ *i*]) is the model’s predicted probability distribution for all 20 amino acids at position *i* conditioned on the structure (*f_nat_*) and the native residues prior in the decoding order (*s_nat_*[≺ *i*]), and *s_nat_*[*i*] is the native residue at position *i*.

Second, we supervised the edges using the negative log composite pseudo-likelihood loss used to train COORDinator (8), shown in Equation 2:

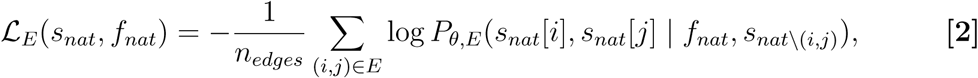

where *n_edges_* is the number of edges in the graph, *i* and *j* are the indices of the residues connected by an edge, and 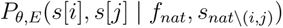 is the model’s predicted probability distribution for all 400 pairs of amino acids at the pair of positions (*i, j*) conditioned on the structure (*f_nat_*) and the native sequence at all other sites (*s_nat_*_\(_*_i,j_*_)_). This distribution is defined according the energies in the Potts model as in Equation 3:

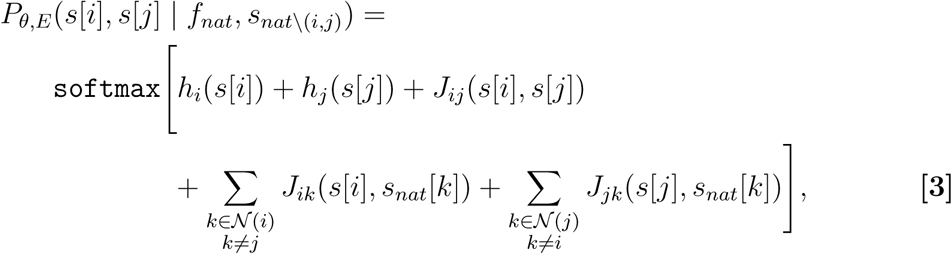

where 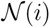 denotes the set of all neighbors of the residue at site *i*. That is, the composite pseudo-likelihood loss rewards the model for assigning a high probability to the native pair of residues at each pair of positions in the sequence, given the native sequence at all other positions.

To train with MSAs, we used the loss function shown in Equation 4:

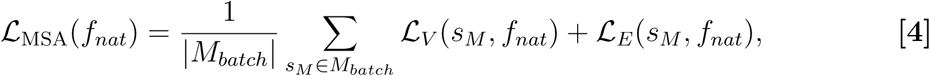

where *M_batch_* is the set of sequences in the filtered MSA subsampled to fit in memory for the current training batch (i.e., such that the total number of residues does not exceed 10,000). We trained a single-site version of PottsMPNN called PottsSingleMPNN that extracts single-site residue probabilities from the conditional distributions of the Potts model as shown in Equation 5:

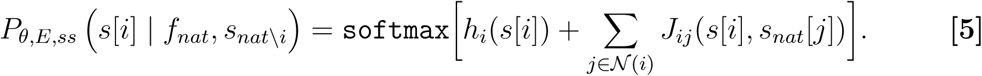

That is, the conditional probability distribution at site *i* is defined by summing the energies of each possible residue at site *i* interacting with all its neighbors given the identities of those residues but not given the identity at site *i*. These probabilities were supervised using the negative log likelihood loss function shown in Equation 6:

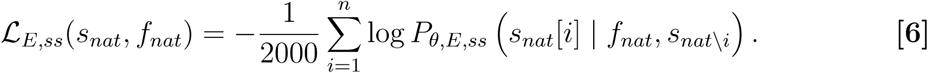

We used the same optimization hyperparameters used to originally train ProteinMPNN (9): an Adam optimizer with *β*_1_ = 0.9, *β*_2_ = 0.98, *∈* = 10^−9^, and the standard attention learning rate schedule with 4000 warm-up steps, and a dropout rate of 10%. We used a batch size of 10,000 tokens. When training without MSAs, we included as many different proteins as possible until the total number of sequence tokens exceeded 10,000. With MSAs, each batch consisted of a single protein, and we included as many randomly sampled sequences from the protein’s MSA as possible given the 10,000 token limit. For each model, we trained for 200 epochs, evaluated the validation loss after each epoch, and tested the model checkpoint with the best validation loss.

### Sequence generation

We used the autoregressive decoding process from ProteinMPNN (9) to iteratively generate a sequence from *P_θ,V_* (*s* | *f*). We used a fixed, randomly chosen decoding order for each protein to control for the effect of the decoding order on model performance by ensuring that every model variant and replicate uses the same order. We used the default ProteinMPNN temperature of 0.1. We experimented with optimizing the autoregressively generated sequences by searching the local space around *s_gen_*. In the local optimization protocol, we iterate over the positions in sequence, evaluate the energy of all amino-acid substitutions at that position according to the Potts model (with the current identity at that position masked), and update the sequence with the best-scoring residue. We iterate in the same decoding order used to generate the sequence. We refer to sequences generated using local optimization with the Potts energies as “optimized.” We experimented with continuing to iterate over the sequence, in the same order used to generate the sequence, until the Potts model finds no favorable substitution at any position (i.e., until convergence). Finally, we experimented with scoring the substitutions using the single-site probabilities. See Energy prediction for details on how energies are calculated.

### Sequence-structure self-consistency

We used two methods for evaluating *P* (*f_nat_* | *s_gen_*). First, we used the monomer ptm version of AlphaFold2 to predict a structure *f_pred_* from *s_gen_* (4). We ran AlphaFold2 without MSAs or templates. We then calculated the TM-score, a metric of structure similarity, between *f_pred_* and *f_nat_* (43). We also examined AlphaFold2 confidence using pLDDT. Second, we used Rosetta to evaluate the compatibility between *s_gen_* and *f_nat_*. To do so, we threaded *s_gen_* onto *f_nat_* and relaxed using the FastRelax protocol with the standard full-atom energy function to minimize steric clashes and optimize side-chain packing (62–64). Following relaxation, the total all-atom energy was evaluated with the same scoring function. To attain the final Rosetta score, we normalize the energy by the protein length. We assessed sequence-structure self-consistency on the CATH 4.2 and PDB-clust test sets. The CATH 4.2 test set consists of 1,108 single-chain proteins, all with fewer than 500 residues. For PDB-clust, to facilitate running AlphaFold2 and Rosetta, we limited the length of proteins to 500 residues, resulting in 702 single- and multi-chain protein structures.

### Energy prediction

To compute energies from single-site probabilities *P_θ,V_* (*s* | *f*)–—from PottsMPNN, ProteinMPNN, or a benchmark model—–we compare the mutant and wild-type probabilities:

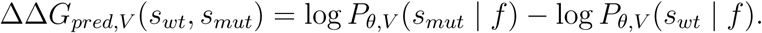

To generate energy predictions from the Potts model, we compute the energy of a mutation as follows:

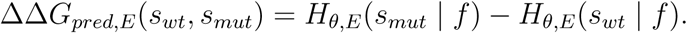

Unless otherwise specified, PottsMPNN energy predictions use the Potts model.

We evaluated energy prediction performance by calculating the Pearson correlation between predicted and observed energies for three datasets. First, we used 238,661 point mutations in the Megascale dataset, which contains stability energy measurements for 298 single-chain proteins (41). Second, we used 2,542 point mutations in the FireProt dataset, which contains stability energy measurements for 88 proteins curated from the literature; energies were measured using various methods (44). Third, we used a dataset of ~4,000 single mutants from a DMS screen over the SARS-CoV-2 receptor binding domain (42). Each mutation has expression data, which we used as a proxy for stability. During training, the models never saw any energy data, but out of an abundance of caution we took steps to prevent data leakage. For the Megascale and FireProt datasets, we removed any proteins that are in the CATH 4.2 or PDB-clust training sets, leaving 202,804 point mutations from 232 proteins for Megascale and 2,301 point mutations from 64 proteins for FireProt. For the SARS-CoV-2 dataset, the structure, 6M0J (65, 66), is in the PDB-clust training set and is not in the CATH 4.2 training set.

## Data, Materials, and Software Availability

The code for PottsMPNN is available at https://github.com/KeatingLab/PottsMPNN. The repository includes training and inference scripts as well as Colab implementations of the sequence design, sequence optimization, and energy prediction tasks discussed in the paper. The repository includes the experimental energy data used to evaluate the models, which are also available at their respective literature sources: Megascale (41), FireProt (44), and SARS-CoV-2 (42).

## Acknowledgments

Research reported in this publication was supported by the National Institute of General Medical Sciences of the National Institutes of Health under R35GM149227 to A.E.K, a Takeda Fellowship awarded to F.B., and a MathWorks Science Fellowship awarded to F.B. The content herein is solely the responsibility of the authors and does not represent the official views of any of the funding organizations. The authors acknowledge the MIT Office of Research Computing and Data for providing high performance computing resources that contributed to the results reported in this paper.

## Supplementary Information

**Fig. S1.**
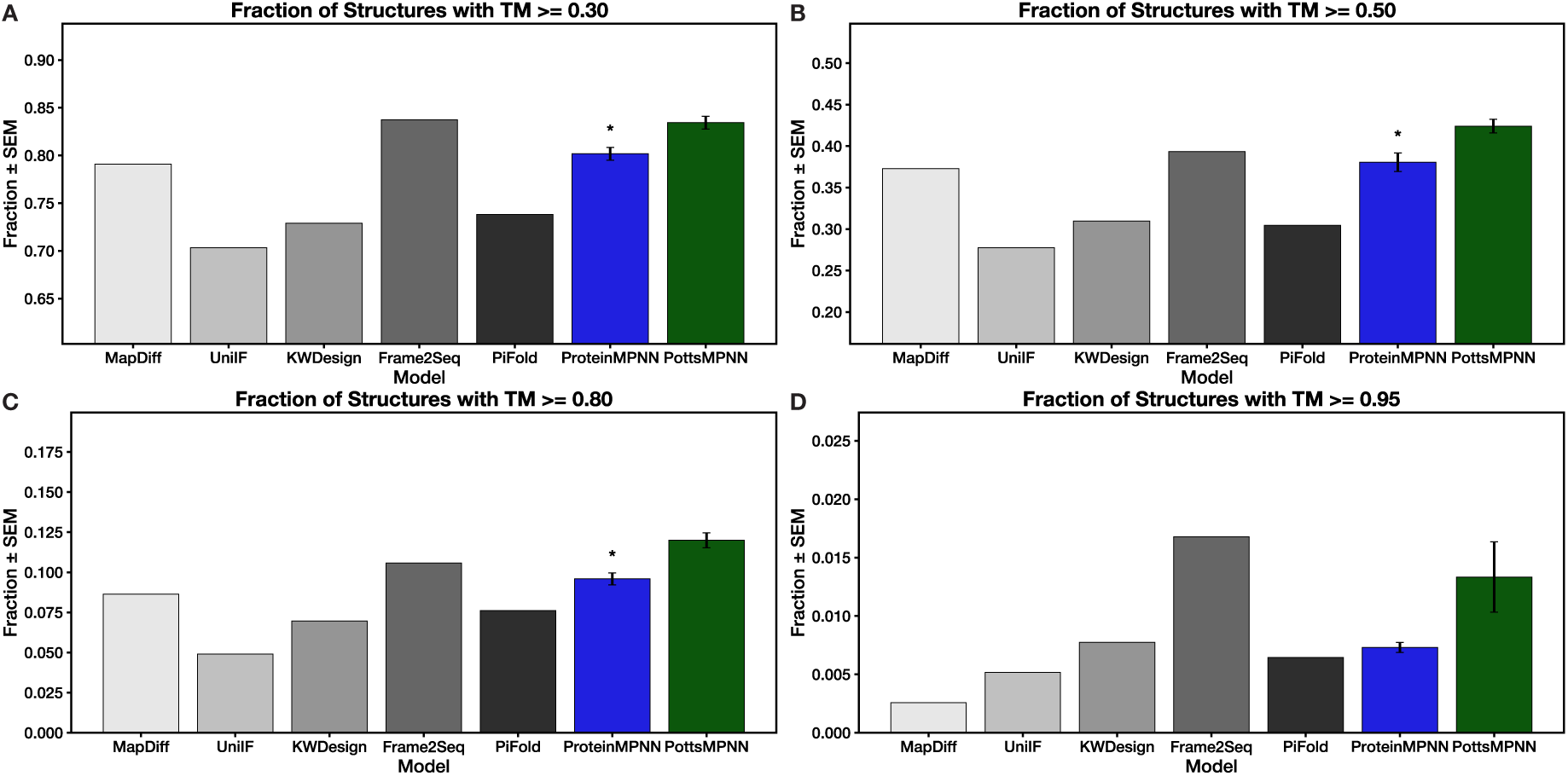
Expanded TM-score results showing the fraction of generated sequences from the CATH 4.2 test set with AlphaFold2 predicted structures that have TM-scores to the native structures above 0.3 (A), 0.5 (B), 0.8 (C), and 0.95 (D). Error bars show SEM results over three retrained model replicates (where available). Stars indicate statistical significance relative to PottsMPNN, assessed using a two-tailed unpaired t-test over average fractions (* p *<* 0.05; no star indicates non-significance).

**Fig. S2.**
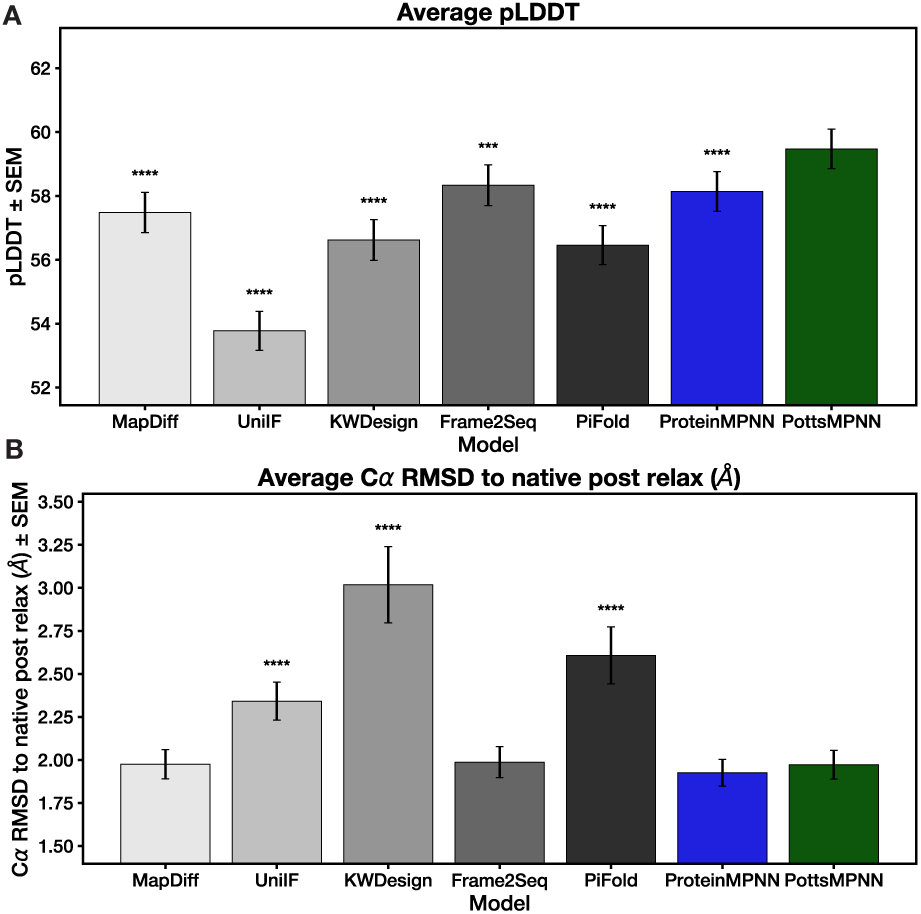
Structure self-consistency confidence results for benchmark models trained and tested on the CATH 4.2 dataset. (A) AlphaFold2 pLDDT scores for structure predictions of sequences generated using each model. (B) C*α* RMSD between the native structures and Rosetta relaxed structures after threading generated sequences onto the native backbone. Error bars show SEM over the proteins in the test set after averaging results over three retrained model replicates. Stars indicate statistical significance relative to PottsMPNN, assessed using a two-tailed paired t-test over per-protein values (* p *<* 0.05, ** p *<* 0.01, *** p *<* 0.001, **** p *<* 0.0001; no star indicates non-significance).

**Fig. S3.**
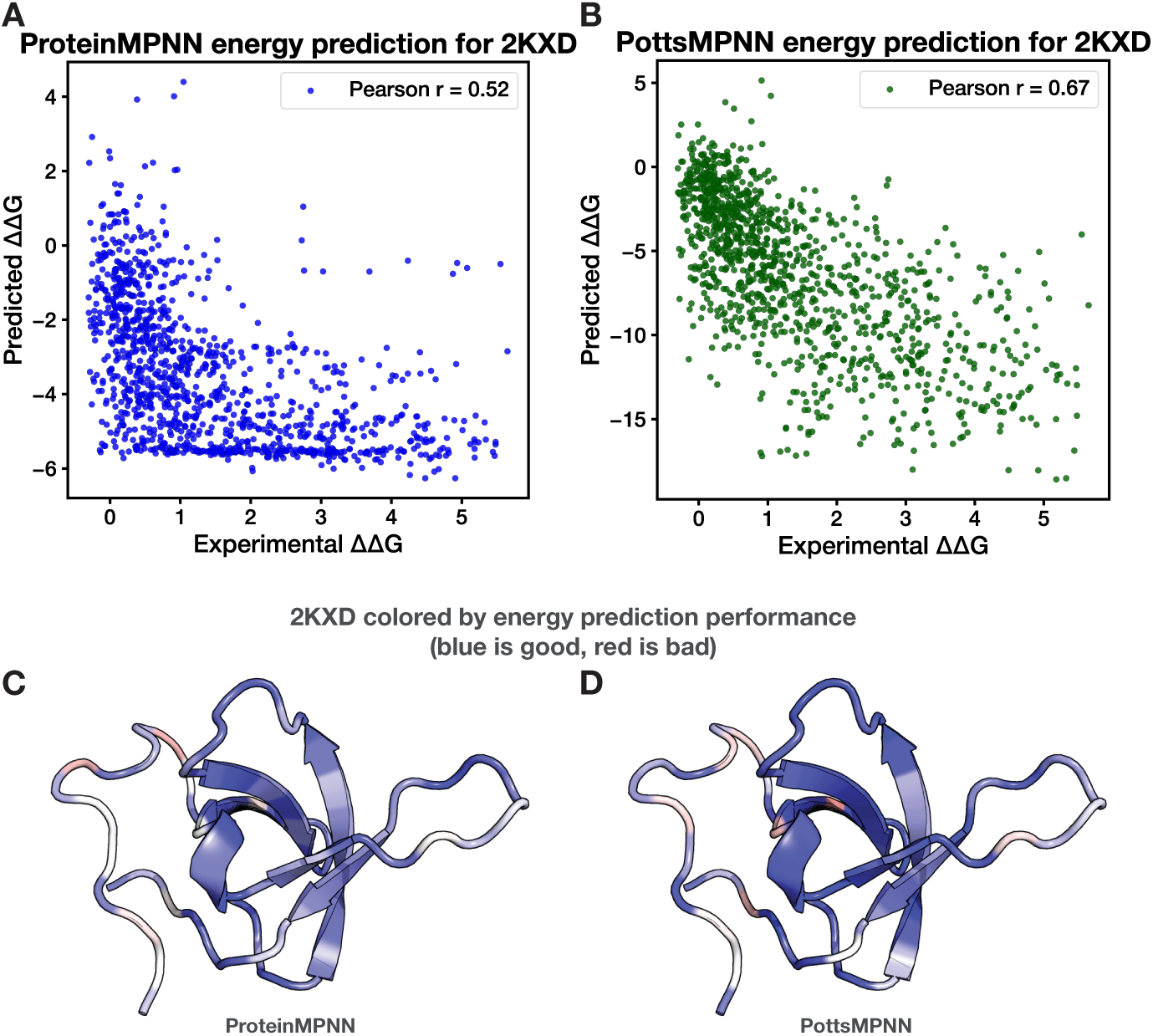
Case study of using ProteinMPNN and PottsMPNN to predict the effect of single-site mutations on protein stability for 2KXD (1, 2), a protein in the Megascale dataset. (A) ProteinMPNN predictions versus experimental ΔΔ*G* values. (B) PottsMPNN predictions versus experimental ΔΔ*G* values. (C) – (D) Heatmaps showing Pearson r coefficients for the correlation between ProteinMPNN predictions (C) and PottsMPNN predictions (D) against experimental ΔΔ*G* values for mutations at individual sites. Blue indicates a positive correlation, and red indicates a negative correlation.

**Fig. S4.**
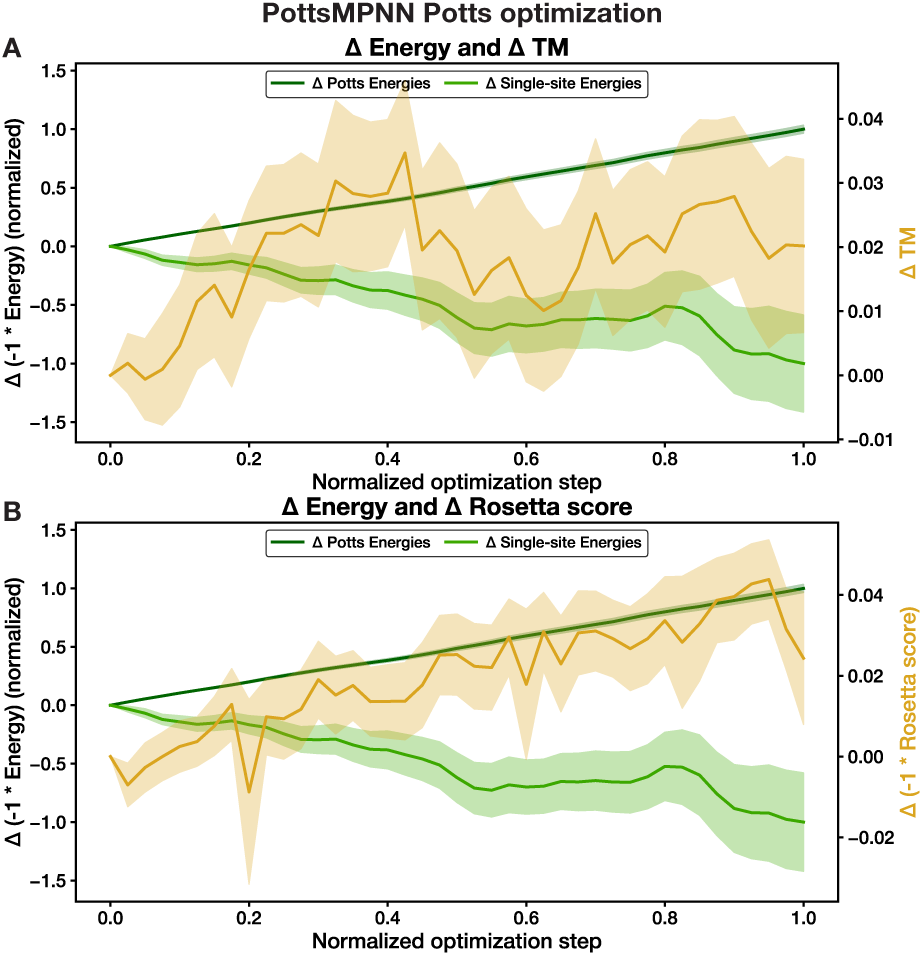
Tracking sequence-structure self-consistency during optimization of PottsMPNN sequences using the Potts model for 100 random sequences designed for structures from the CATH 4.2 test set. (A) The average normalized change in energy as assessed by the Potts model (dark green) and by the single-site energies (light green) compared to the average change in the TM-score between the native structure and the AlphaFold2 predicted structure (yellow). (B) The average normalized change in energy as assessed by the Potts model (dark green) and by the single-site energies (light green) compared to the average change in the Rosetta score after threading generated sequences onto the native backbone and relaxing (yellow). All scores are plotted such that a higher value is more desirable. Shaded regions indicate the SEM.

**Fig. S5.**
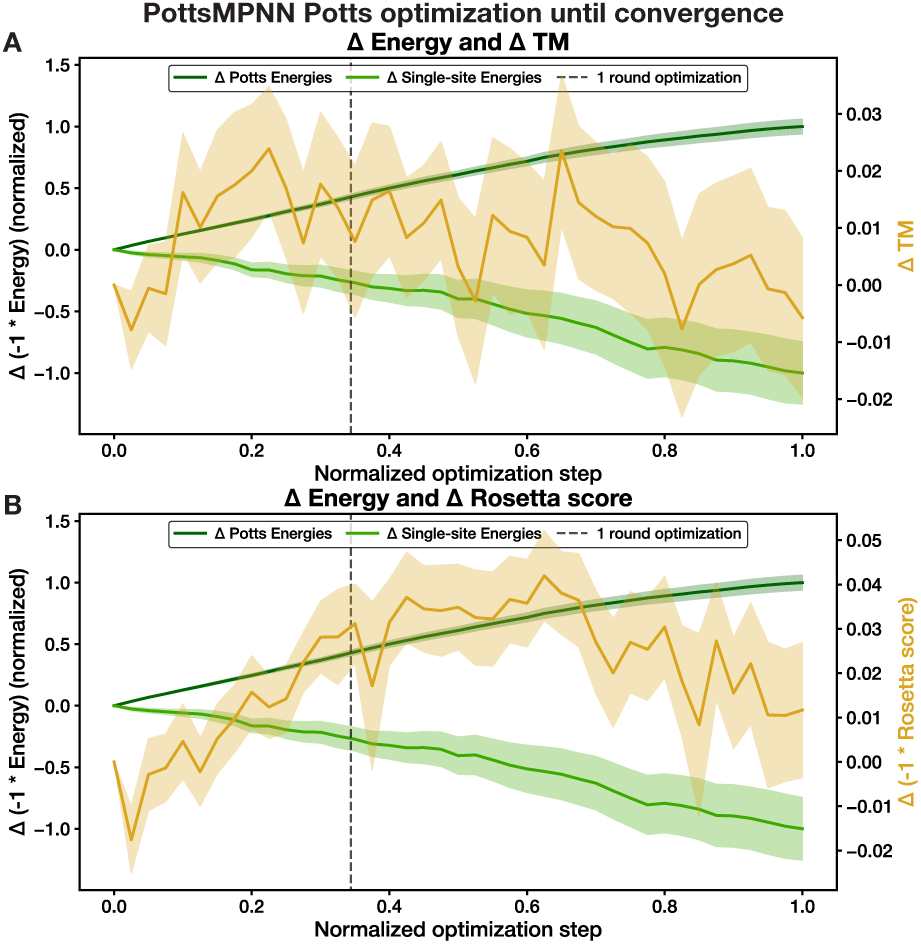
Tracking sequence-structure self-consistency during optimization of PottsMPNN sequences using the Potts model for 100 random sequences designed for structures from the CATH 4.2 test set until no more favorable mutations are found. (A) The average normalized change in energy as assessed by the Potts model (dark green) and by the single-site energies (light green) compared to the average change in the TM-score between the native structure and the AlphaFold2 predicted structure (yellow). (B) The average normalized change in energy as assessed by the Potts model (dark green) and by the single-site energies (light green) compared to the average change in the Rosetta score after threading generated sequences onto the native backbone and relaxing (yellow). All scores are plotted such that a higher value is more desirable. Shaded regions indicate the SEM. The dotted line indicates the average step at which the standard optimization would have stopped (i.e., after each position in the sequence has been visited once).

**Fig. S6.**
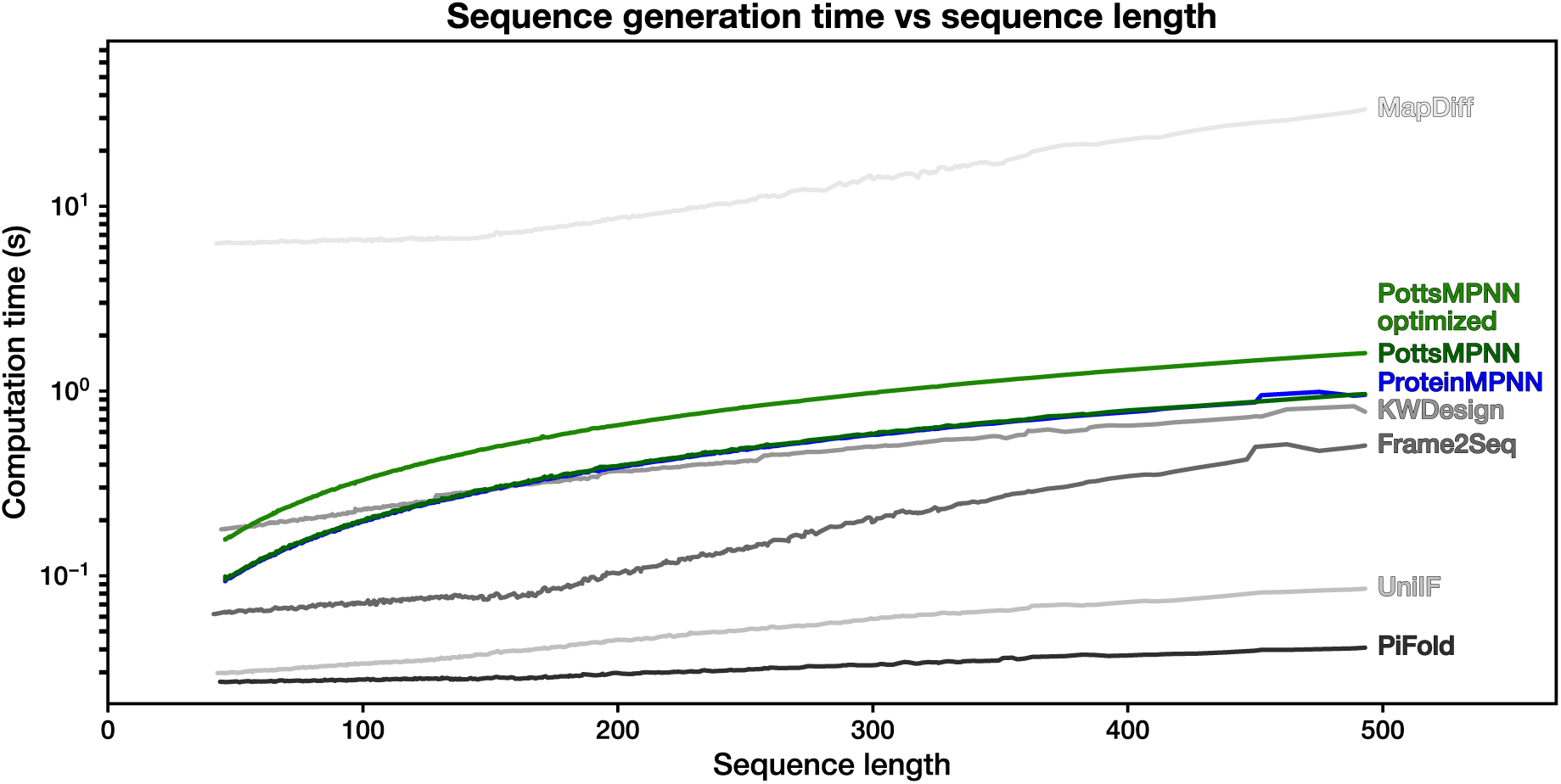
Time needed to generate 1 sequence for each protein in the CATH 4.2 test set when running the models on an Nvidia A6000 GPU, plotted by sequence length.

**Fig. S7.**
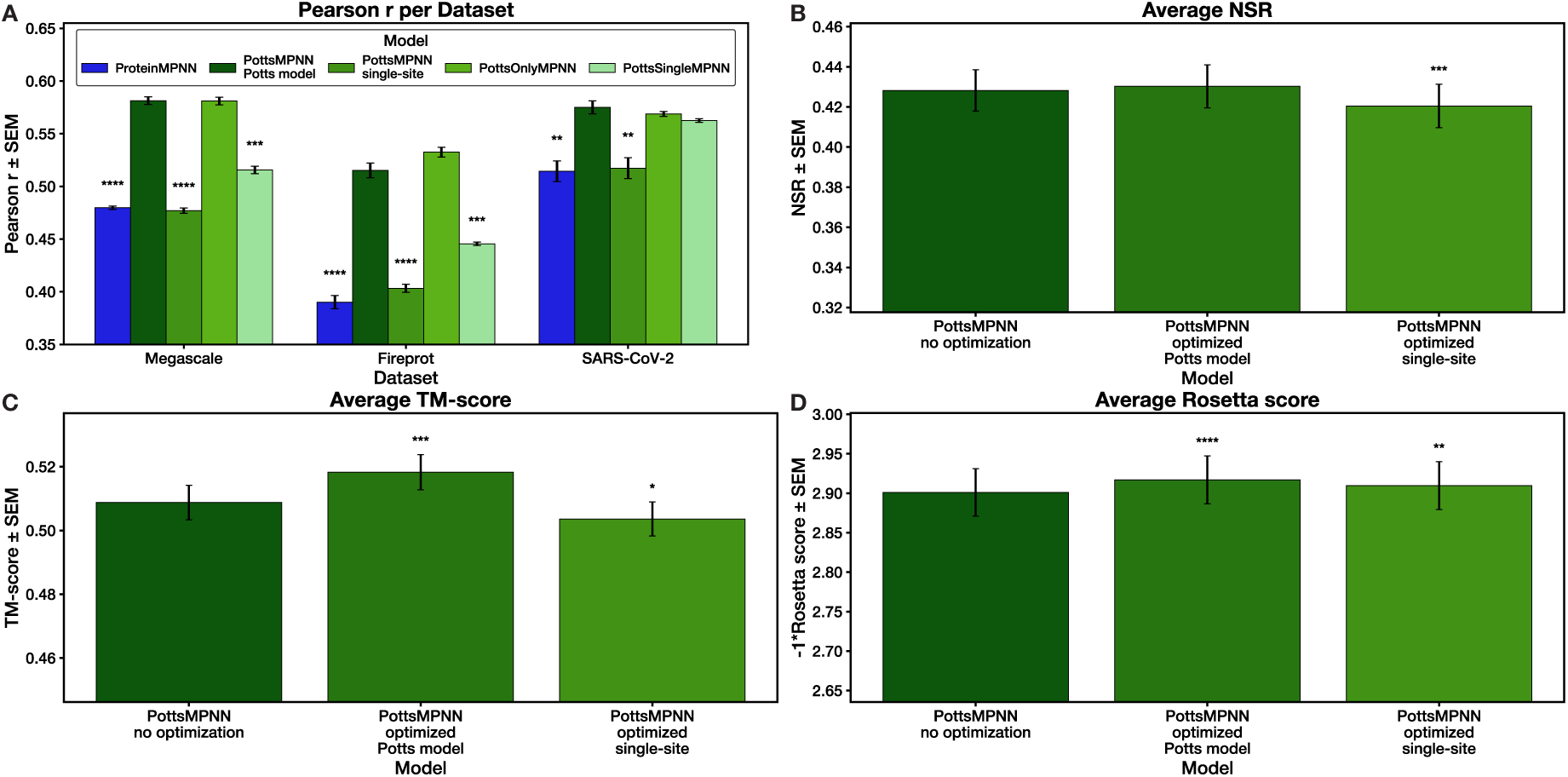
PottsMPNN Potts model experimentation results for models trained and tested on CATH 4.2. (A) Pearson r for using ProteinMPNN, the Potts model from PottsMPNN, the single-site probabilities from PottsMPNN, PottsOnlyMPNN, and PottsSingleMPNN to predict the effect of single-site mutations on protein stability for three datasets. (B) NSR results for base PottsMPNN (i.e., without optimization), PottsMPNN with one pass of optimization using the Potts model, and PottsMPNN with one pass of optimization using the single-site probabilities. (C) TM-scores between native structures and AlphaFold2 structures predicted for sequences generated using each optimization method. (D) Rosetta scores after threading generated sequences onto the native backbone and relaxing for each optimization method. For (A), error bars show SEM over at least two retrained model replicates. For (B) – (D), error bars show SEM over the proteins in the test set after averaging results over at least two retrained model replicates. Stars indicate statistical significance relative to PottsMPNN Potts model for (A) and to PottsMPNN no optimization for (B) – (D), assessed using a two-tailed unpaired t-test over average Pearson r values for (A) and a two-tailed paired t-test over per-protein values for (B) – (D) and (* p *<* 0.05, ** p *<* 0.01, *** p *<* 0.001, **** p *<* 0.0001; no star indicates non-significance).

**Fig. S8.**
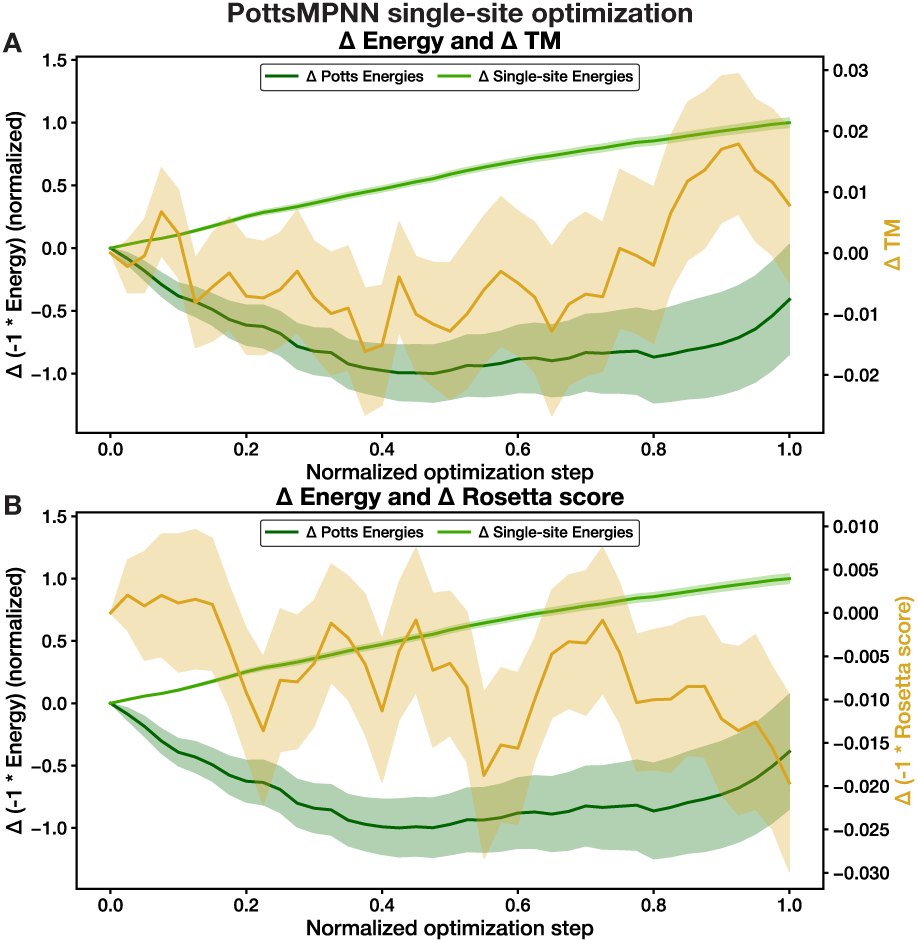
Tracking sequence-structure self-consistency during optimization of PottsMPNN sequences using the single-site energies for 100 random sequences designed for structures from the CATH 4.2 test. (A) The average normalized change in energy as assessed by the Potts model (dark green) and by the single-site energies (light green) compared to the average change in the TM-score between the native structure and the AlphaFold2 predicted structure (yellow). (B) The average normalized change in energy as assessed by the Potts model (dark green) and by the single-site energies (light green) compared to the average change in the Rosetta score after threading generated sequences onto the native backbone and relaxing (yellow). All scores are plotted such that a higher value is more desirable. Shaded regions indicate the SEM.

**Fig. S9.**
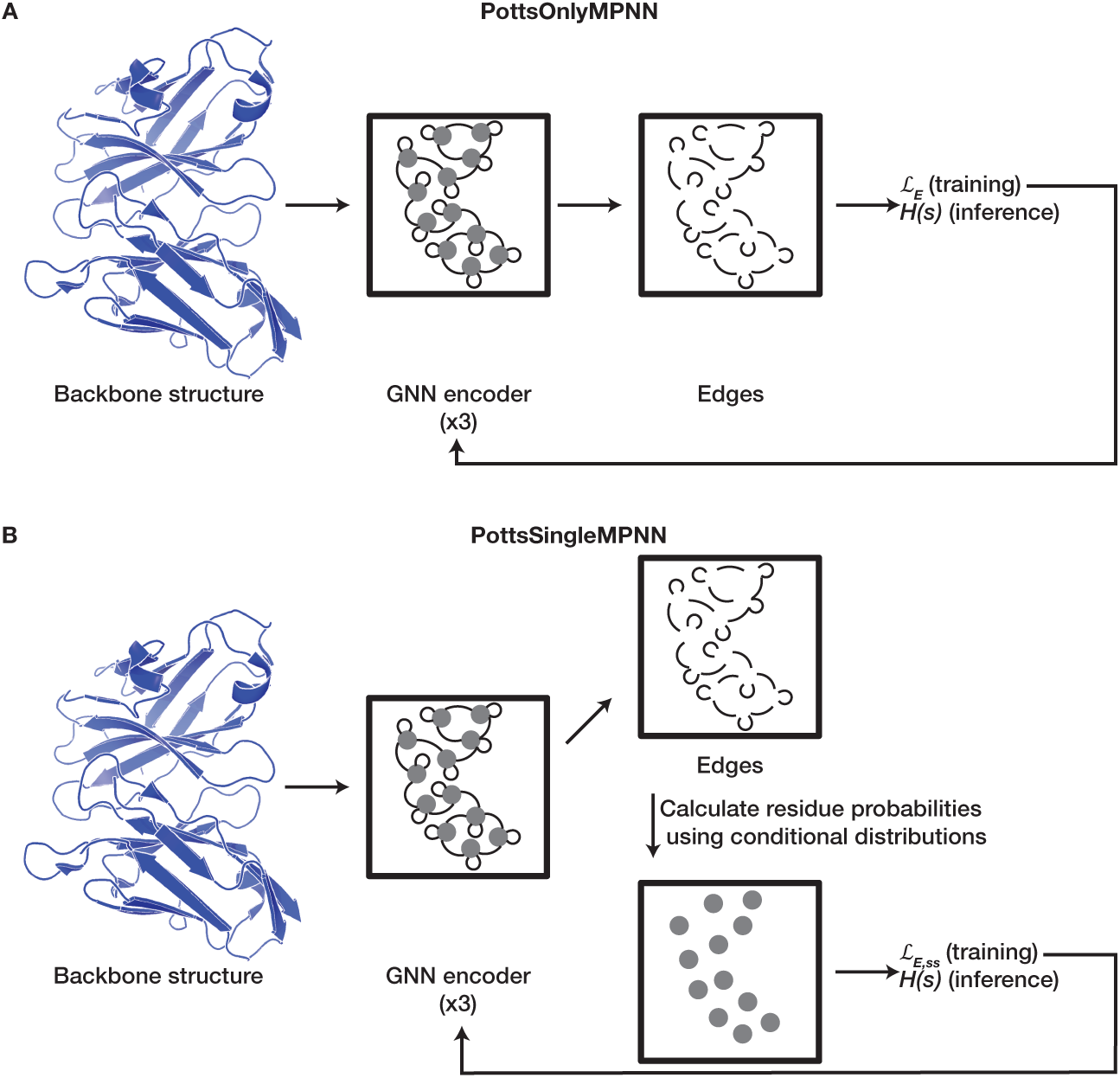
Overview of the PottsMPNN ablated architectures. (A) Overview of the PottsOnlyMPNN architecture. The graph edges, including self-edges, are supervised to learn single and pairwise residue interaction energies in the form of a Potts model (*H*(*s*)) that is used to compute structure energies. The Potts model is supervised using a negative log composite pseudo-likelihood loss (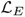) to maximize the probability of native residue pairs. (B) Overview of the PottsSingleMPNN architecture. From the final edge embeddings, conditional probabilities are calculated to generate single-site amino-acid probability distributions, which are supervised by a single-site negative log-likelihood loss (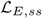); during inference, the Potts model (*H*(*s*)) is used to compute structure energies.

**Fig. S10.**
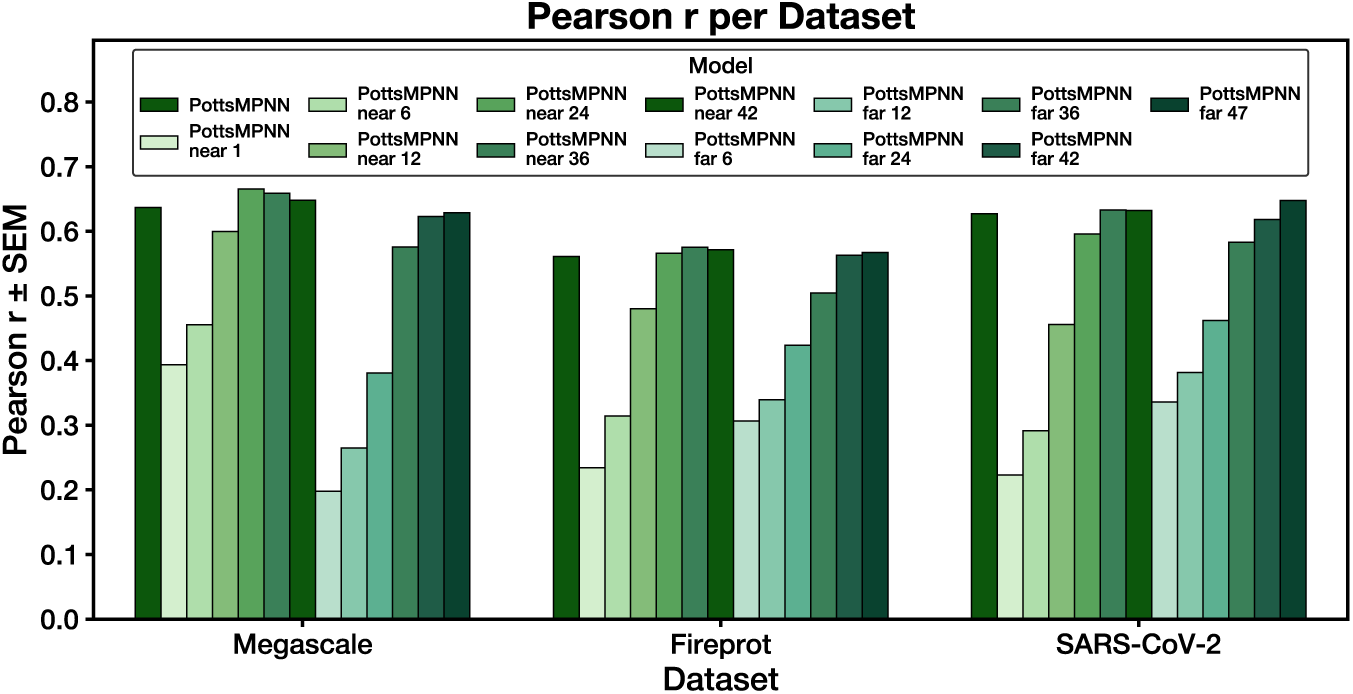
The impact of including different residue-residue interactions on the Pearson r values for predicting the effect of single-site mutations. Each bar represents the Pearson r resulting from using pair-energies in the Potts model for the indicated neighbor set (e.g., near 6 indicates only the pair-energies with the 6 closest neighbors for each residue were used; far 6 indicates only the pair-energies with the 6 furthest neighbors–—out of the 48-nearest neighbors–—for each residue were used). Models were trained on PDB-clust.

**Fig. S11.**
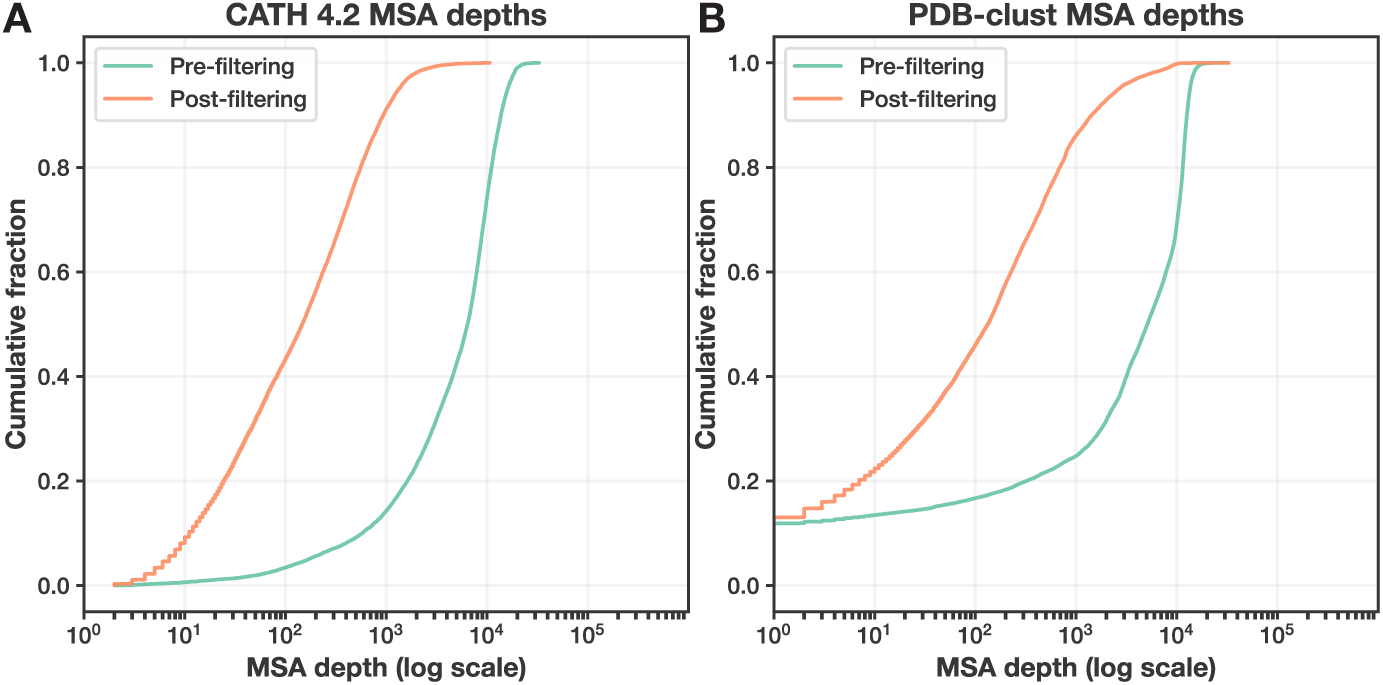
MSA depths pre- and post-filtering for CATH 4.2 MSAs (A) and PDB-clust MSAs (B). MSAs were filtered using a sequence identity minimum of 50%, a deletion percentage maximum of 20%, and an insertion percentage maximum of 20%. The *~*17% of proteins in the PDB-clust dataset with MSAs of depth 1 correspond to multi-chain proteins for which the paired MSAs are empty.

**Fig. S12.**
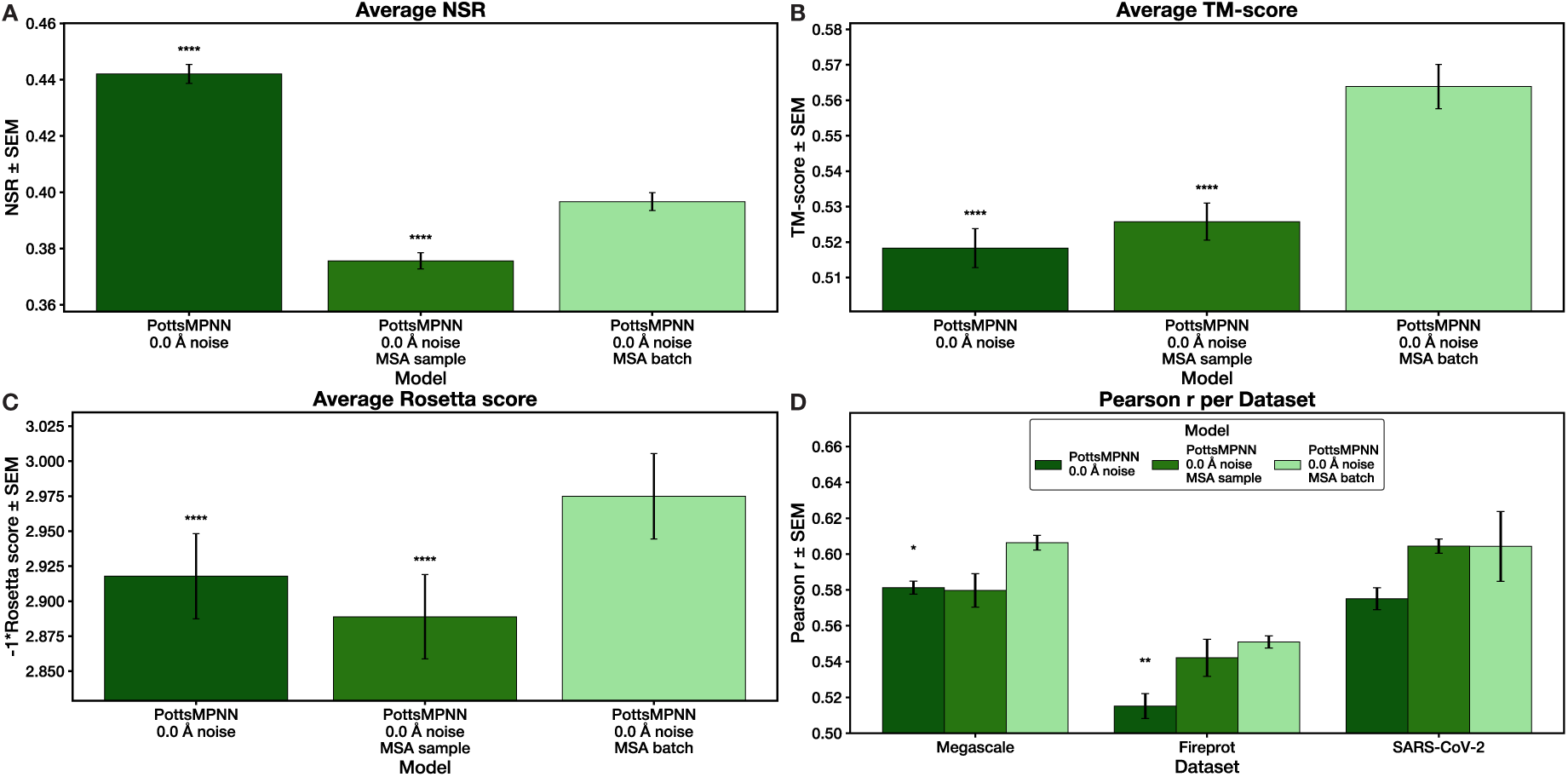
Using multiple sequences from the MSA during each iteration (MSA batch) versus sampling a single sequence (MSA sample) is key to the improved performance from training with an MSA loss on the CATH 4.2 dataset. (A) NSR results for each model. (B) TM-scores between native structures and AlphaFold2 predicted structures for sequences generated using each model. (C) Rosetta scores after modeling generated sequences on the native backbones for each model. (D) Pearson r for using each model to predict the effect of single-site mutations on protein stability for three datasets. For (A) – (C), error bars show SEM over the proteins in the test set after averaging results over at least two retrained model replicates; for (D), error bars show SEM over at least two retrained model replicates. Stars indicate statistical significance relative to PottsMPNN 0.0 Å noise MSA batch, assessed using a two-tailed paired t-test over per-protein values for (A)–(C) and a two-tailed unpaired t-test over average Pearson r values for (D) (* p *<* 0.05, ** p *<* 0.01, *** p *<* 0.001, **** p *<* 0.0001; no star indicates non-significance).

**Fig. S13.**
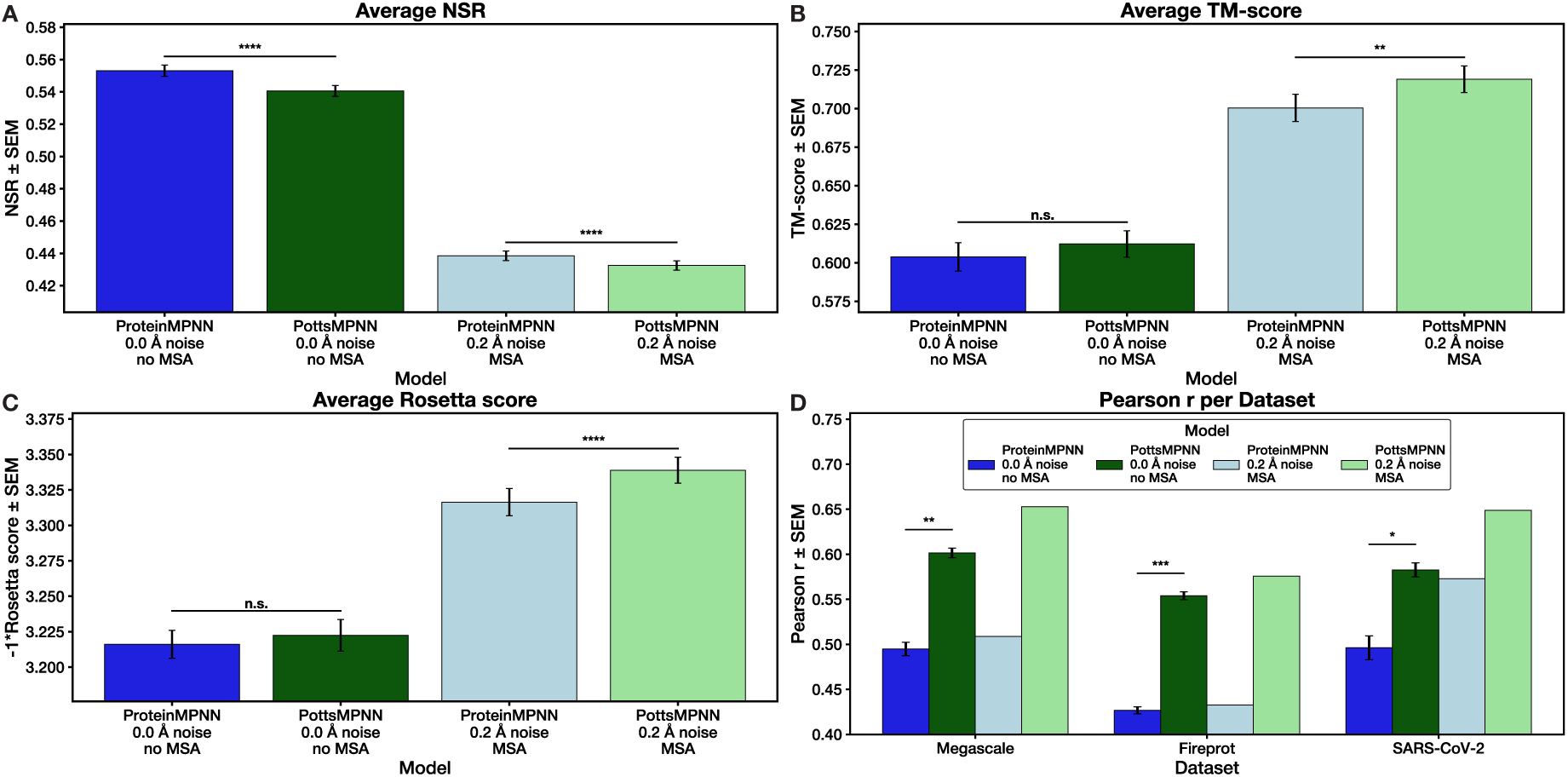
Training with an MSA loss function on the PDB-clust dataset improves performance. (A) NSR results for each model. (B) TM-scores between native structures and AlphaFold2 predicted structures for sequences generated using each model. (C) Rosetta scores after threading generated sequences onto the native backbone and relaxing for each model. (D) Pearson r for using each model to predict the effect of single-site mutations on protein stability for three datasets. For (A) – (C), error bars show SEM over the proteins in the test set after averaging results over at least two retrained model replicates for the no MSA models; for (D), error bars show SEM over at least two retrained model replicates for the no MSA models. Only one replicate was trained for the MSA models. Stars indicate statistical significance comparing ProteinMPNN to PottsMPNN for each condition, assessed using a two-tailed paired t-test over per-protein values for (A) – (C) and a two-tailed unpaired t-test over average Pearson r values for (D) (* p *<* 0.05, ** p *<* 0.01, *** p *<* 0.001, **** p *<* 0.0001; n.s. indicates non-significance; no bar indicates no statistical test was performed).

**Fig. S14.**
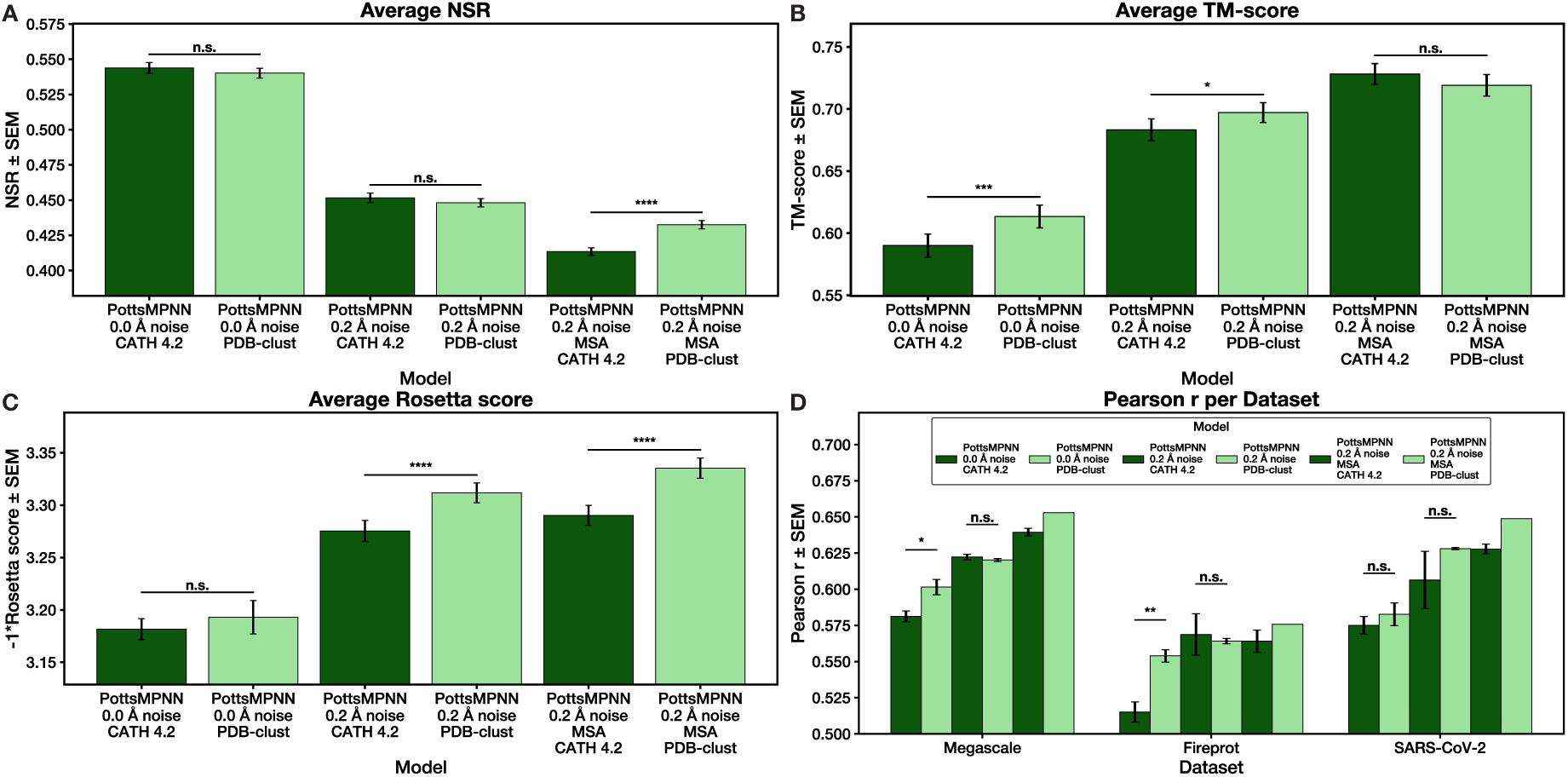
Training on the PDB-clust dataset is superior to training on the CATH 4.2 dataset when evaluated on a subset of the PDB-clust dataset that is non-overlapping with the CATH 4.2 train set. (A) NSR results for each model. (B) TM-scores between native structures and AlphaFold2 predicted structures for sequences generated using each model. (C) Rosetta scores after modeling generated sequences onto the native backbones. (D) Pearson r for using each model to predict the effect of single-site mutations on protein stability for three datasets. For (A) – (C), error bars show SEM over the proteins in the test set after averaging results over at least two retrained model replicates; for (D), error bars show SEM over at least two retrained model replicates (only 1 replicate of PottsMPNN 0.2 Å noise MSA model was trained on PDB-clust). Stars indicate statistical significance comparing training on CATH 4.2 to training on PDB-clust for each condition, assessed using a two-tailed paired t-test over per-protein values for (A) – (C) and a two-tailed unpaired t-test over average Pearson r values for (D) (* p *<* 0.05, ** p *<* 0.01, *** p *<* 0.001, **** p *<* 0.0001; n.s. indicates non-significance; no bar indicates no statistical test was performed).

**Fig. S15.**
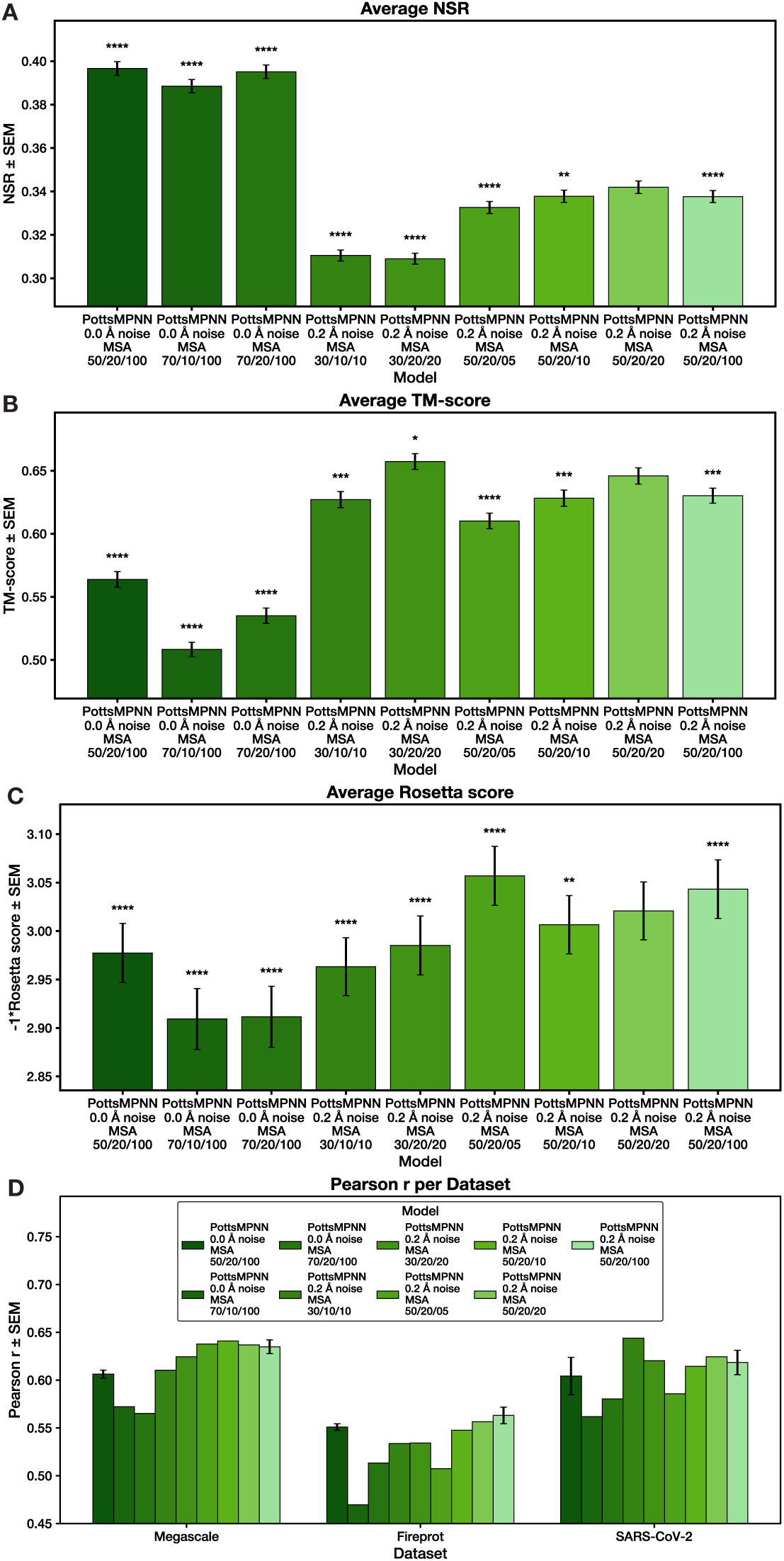
MSA filtering hyperparameter experiments. (A) NSR results for each model. The hyperparameters are the last line of each label. The first number represents the sequence identity minimum; the second number represents the deletion percentage maximum; and the third number represents the insertion percentage maximum. (B) TM-scores between native structures and AlphaFold2 predicted structures for sequences generated using each model. (C) Rosetta scores after threading generated sequences onto the native backbone and relaxing for each model. (D) Pearson r for using each model to predict the effect of single-site mutations on protein stability for three datasets. For (A) – (C), error bars show SEM over the proteins in the CATH 4.2 test set after averaging over two model replicates; for (D), error bars show SEM over two model replicates. For (D), lack of error bars indicates only one model replicate was trained. Stars indicate statistical significance relative to PottsMPNN 0.2 Å noise MSA 50/20/20, assessed using a two-tailed paired t-test over per-protein values for (A) – (C) and a two-tailed unpaired t-test over average Pearson r values for (D) (* p *<* 0.05, ** p *<* 0.01, *** p *<* 0.001, **** p *<* 0.0001; no star indicates non-significance).

## Notes

https://github.com/KeatingLab/PottsMPNN/

### Competing Interest Statement

The authors have declared no competing interest.

### Summary of Updates

Added additional references on the use of learned pairwise-decomposable energy functions for protein sequence design.

